# ITPK1 is an InsP_6_/ADP phosphotransferase that controls systemic phosphate homeostasis in Arabidopsis

**DOI:** 10.1101/2020.05.18.100297

**Authors:** Esther Riemer, Debabrata Laha, Robert K. Harmel, Philipp Gaugler, Verena Pries, Michael Frei, Mohammad-Reza Hajirezaei, Nargis P. Laha, Lukas Krusenbaum, Robin Schneider, Henning J. Jessen, Adolfo Saiardi, Dorothea Fiedler, Gabriel Schaaf, Ricardo F.H. Giehl

**Affiliations:** Department of Plant Nutrition, Institute of Crop Science and Resource Conservation, Rheinische Friedrich-Wilhelms-Universität Bonn, 53115 Bonn, Germany; Medical Research Council Laboratory for Molecular Cell Biology (MRC-LMCB), University College London, London WC1E 6BT, United Kingdom; Leibniz-Forschungsinstitut für Molekulare Pharmakologie, 13125 Berlin, Germany and Department of Chemistry, Humboldt Universität zu Berlin, 12489 Berlin, Germany; Department of Physiology & Cell Biology, Leibniz-Institute of Plant Genetics and Crop Plant Research, 06466 Gatersleben, Germany; Department of Chemistry and Pharmacy and CIBSS-Centre for Integrative Biological Signalling Studies, Albert-Ludwigs University Freiburg, 79104 Freiburg, Germany

## Abstract

In plants, phosphate (P_i_) homeostasis is regulated by the interaction of P_i_ starvation response transcription factors (PHRs) with stand-alone SPX proteins, which act as sensors for inositol pyrophosphates (PP-InsPs). Recently, ITPK1 was shown to generate the PP-InsP InsP_7_ from InsP_6_ *in vitro*, but the importance of this activity in P_i_ signaling remained unknown. Here, we show that uncontrolled P_i_ accumulation in ITPK1-deficient plants is accompanied by impaired P_i_-dependent InsP_7_ and InsP_8_ synthesis. Reciprocal grafting demonstrates that P_i_ starvation responses are mainly controlled by ITPK1 activity in shoots. Nuclear magnetic resonance assays and PAGE analyses with recombinant protein reveal that besides InsP_6_ phosphorylation, ITPK1 is also able to catalyze ATP synthesis using 5-InsP_7_ but not any other InsP_7_ isomer as a P-donor when ATP is low. Additionally, we show that the dynamic changes in InsP_7_ and InsP_8_ to cellular P_i_ are conserved from land plant species to human cells, suggesting that P_i_-dependent PP-InsP synthesis is a common component of P_i_ signaling across kingdoms. Together, our study demonstrates how P_i_-dependent changes in nutritional and energetic states modulate ITPK1 activities to fine-tune the synthesis of PP-InsPs.

## INTRODUCTION

In order to maintain cellular P_i_ homeostasis, plants have evolved complex sensing and signaling mechanisms that adjust whole-plant P_i_ demand with external P_i_ availability. Although many molecular players involved in these responses have been identified, the exact mechanism of P_i_ sensing in complex organisms, such as plants, still remains largely unknown. In the model species *Arabidopsis thaliana*, the MYB transcription factors PHOSPHATE STARVATION RESPONSE 1 (PHR1) and its closest paralog PHR1-LIKE1 (PHL1) control the expression of the majority of P_i_ starvation-induced (PSI) genes and influence numerous metabolic and developmental adaptations induced by P_i_ deficiency^1, 2^. In agreement with their regulatory function, a subset of P_i_ deficiency-induced genes is deregulated in *phr1 phl1* mutants. However, since *PHR1* itself is not transcriptionally regulated by P_i_ deficiency, the existence of a post-translational control of PHR1 has been proposed^1^. Emerging evidence indicates that a class of stand-alone SPX proteins negatively regulate the activity of PHR transcription factors in different plant species^3, 4, 5, 6, 7, 8^. According to these studies, when plants have access to sufficient P_i_, SPX proteins bind with high affinity to PHRs, thereby preventing binding of these transcription factors to DNA. Under low P_i_, the affinity of SPX proteins towards PHRs is decreased, thus allowing these transcription factors to activate their transcriptional targets^4^.

The *in vivo* interaction of PHR1 and SPX1 is influenced by P_i_^3, 4, 5^, suggesting that this mechanism could represent a direct link between P_i_ perception and downstream signaling events. However, the dissociation constants for P_i_ itself in a SPX-PHR complex ranged from 10 mM to 20 mM^3, 4, 5^, while P_i_ levels of only 60 μM to 80 μM have been recorded in the cytosol of plant cells^9^. A later study, demonstrated that SPX domains can actually act as receptors for inositol pyrophosphates (PP-InsPs) and isothermal titration calorimetry experiments revealed that 5PP-InsP_5_ (hereafter referred to as 5-InsP_7_) interacted more strongly with SPX domains than P_i_^10^. In these assays, interaction of rice OsPHR2 and OsSPX4 was promoted at 5-InsP_7_ concentrations as low as 20 μM^10^. Although the isomeric nature of plant InsP_7_ in vegetative tissues remains unknown, it has been proposed that the activity of PHRs is regulated by direct interaction with PP-InsPs. In support of this hypothesis, an Arabidopsis mutant for *INOSITOL PENTAKISPHOSPHATE 2-KINASE* (*IPK1*), which is compromised in the synthesis of the PP-InsP precursor inositol hexakisphosphate (InsP_6_), exhibits constitutive PSR and increased P_i_ accumulation when grown on sufficient P_i_ availability^11, 12^. More recently, other mutants with compromised synthesis of PP-InsPs have been shown to exhibit disturbed PSR and P_i_ over-accumulation phenotypes^13, 14, 15^, and it was observed that the levels of different inositol polyphosphates (InsPs) are significantly altered in P_i_-deficient plants^13, 14^. Interestingly, polyacrylamide gel electrophoresis (PAGE) analyses revealed that InsP_8_ levels increase in P_i_-sufficient plants and decrease as plants are exposed to P_i_ deficiency^14^, suggesting that the enzymes involved in the synthesis of PP-InsPs could act as regulators of P_i_ homeostasis in plants. This regulation was first discovered in the yeast *Saccharomyces cerevisiae* in which cellular levels of PP-InsPs decrease when exposed to P_i_-deficient medium^10, 16^. However, the biosynthetic pathway resulting in P_i_-dependent PP-InsP synthesis still remains largely unresolved and it also remains unclear if this response is conserved in the plant lineage and across kingdoms.

In plants, synthesis of InsP_8_ is mediated by VIH1 and VIH2^17^, a class of bifunctional kinase/phosphatase enzymes^15^ sharing homology to the yeast and animal Vip1/PPIP5Ks^17, 18,19^. Although *vih1* and *vih2* single mutants do not exhibit impaired P_i_ accumulation^13^, deletion of both VIHs results in severe growth defects and P_i_ overaccumulation^14, 15^. Since plant genomes do not encode homologues of the metazoan and yeast InsP_6_ kinases IP6K/Kcs1, it has since long remained elusive how plants synthesize InsP_7_. Using a yeast complementation assay, we recently identified Arabidopsis ITPK1 and ITPK2 as novel plant InsP_6_ kinases and demonstrated that ITPK1 generates the symmetric InsP_7_ isomer 5-InsP_7_, the major form identified in seed extracts^20^. More recently, we further showed that InsP_7_ and InsP_8_ levels are compromised in TPK1-deficient plants, demonstrating that ITPK1 functions as an InsP_6_ kinase *in planta*^21^. Considering that InsP_7_ is the precursor for InsP_8_ synthesis, the next challenge is to determine how InsP_7_ levels respond to the plant’s P_i_ status. A recent study reported that InsP_7_ levels increase in shoots of P_i_-deficient Arabidopsis plants as determined by HPLC analysis of [^32^P]-P_i_-labeled plant extracts^13^. However, this response is opposite to what has been described in yeast^16^ and mammalian cells^22^ and is inconsistent with increased InsP_8_ levels detected by PAGE in P_i_-sufficient Arabidopsis plants^14^. Considering that^32^P-labeling entails complicated molecule assignment and does not provide a mass assay of the inositol backbone but a readout for pyrophosphate moiety turnover, these results await confirmation via alternative approaches. Importantly, it remains unclear whether ITPK1 and ITPK2 contribute to PP-InsP synthesis in a P_i_- and/or cellular energy status-dependent manner and how the proposed regulatory activity of InsP_8_ to suppress PSR might be deactivated once plants experience P_i_ deficiency.

Here, we combine strong anion exchange chromatography (SAX)-HPLC analyses of [^3^H]-inositol-labeled seedlings and PAGE to investigate P_i_-dependent changes in the levels of InsP_6_, InsP_7_ and InsP_8_ across diverse species and several *Arabidopsis* mutants. We demonstrate that specifically in shoots, PP-InsPs decrease during P_i_ starvation and strongly increase after P_i_ resupply. P_i_-dependent regulation of PP-InsPs was also highly conserved from diverse plant species to human cells. We find that ITPK1 is critical for the synthesis of PP-InsPs involved in the regulation of P_i_ homeostasis. In addition, we demonstrate that ITPK1-mediated conversion of InsP_6_ to 5-InsP_7_ requires high ATP concentrations and that ITPK1 has ADP phosphotransferase activity under conditions of decreased adenylate energy charge. These results provide a further link between P_i_-dependent changes in nutritional and energetic states with the synthesis of regulatory PP-InsPs. Finally, our study demonstrates that ITPK1 activity in shoots regulates PSR responses in a PHR1/PHL1-dependent manner, revealing that ITPK1 is a critical component of the systemic P_i_ sensing and signaling mechanism in plants.

## RESULTS

### Loss of ITPK1 but not ITPK2 affects P_i_ homeostasis in Arabidopsis

Previously, it has been reported that shoot P_i_ concentrations are significantly changed in a number of InsP biosynthesis mutants grown in hydroponics^13^. We assessed the growth of several of these mutants on soil and compared total P_i_ levels in shoots (measured with ICP-OES) with that of WT and the P_i_-overaccumulator *pho2-1*. We confirmed that *ipk1-1* and *itpk1* plants accumulated significantly more total P_i_ in shoots, reaching levels comparable to those detected in *pho2-1* (Fig. 1a and 1b). To a much lesser degree, shoot P_i_ levels were also significantly increased in shoots of *mips1* and *vih2-4* plants, whereas P_i_ concentrations in all other mutants assessed, including *itpk2-2*, were similar to their respective WTs. A full elemental analysis indicated that the concentrations of other nutrients were largely unaffected in shoots of *itpk1* plants as compared to WT (Suppl. Fig. 1). Total P_i_ levels were also significantly increased in flowers and seeds of *itpk1* plants, although the relative changes were less marked as those detected in rosette leaves (Suppl. Fig. 2). Excessive P_i_ accumulation could be almost completely complemented in transgenic lines expressing the genomic *ITPK1* fragment in the *itpk1* background (Fig. 1c and 1d), showing that P_i_ overaccumulation phenotype was indeed associated with the loss of ITPK1.

**Fig. 1.**
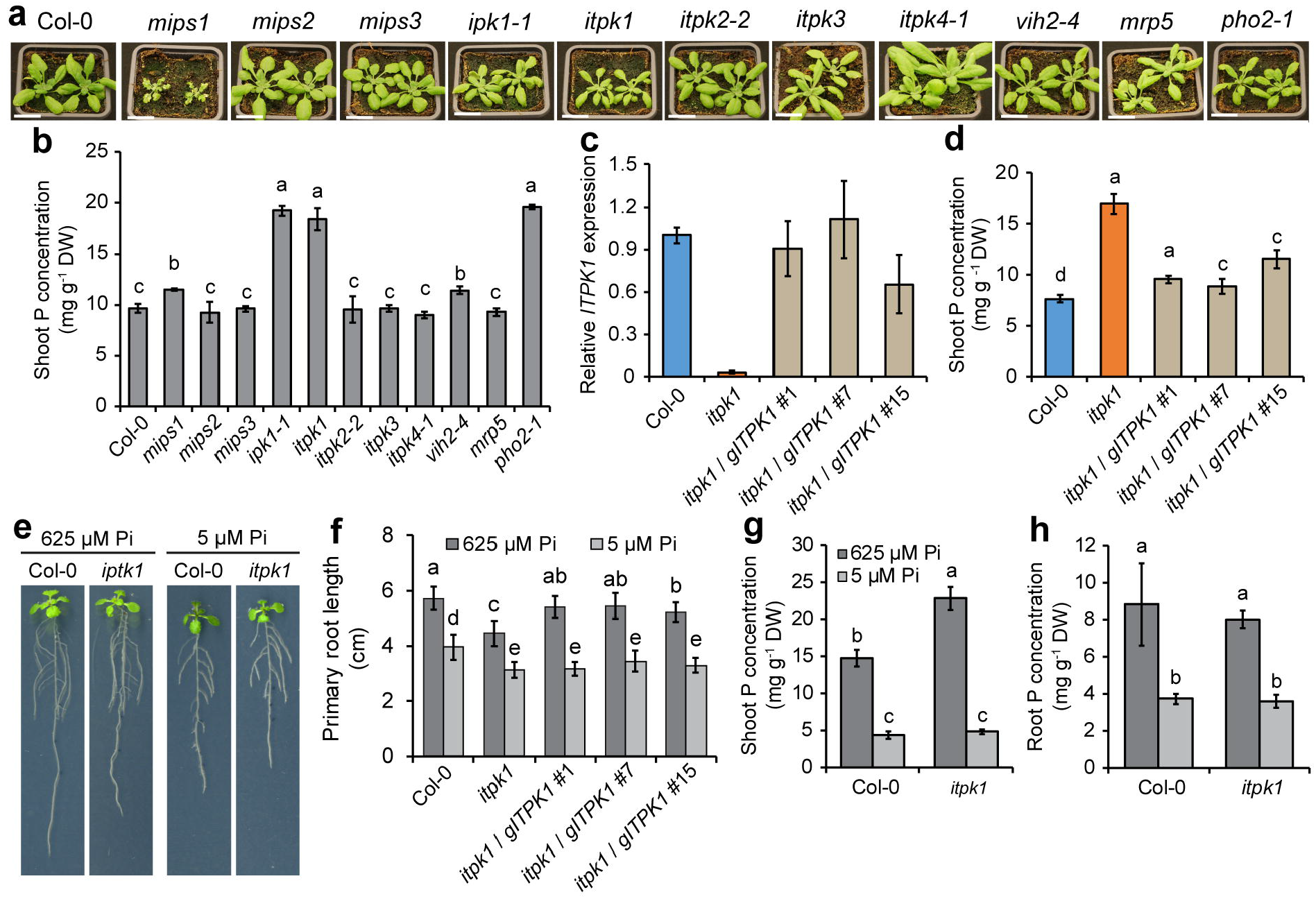
Role of distinct InsP kinases in plant P_i_ accumulation. **a-b** Photographs of 3-week-old *Arabidopsis* plants grown on peat-substrate (**a**), and total P_i_ levels in shoots (**b**) of WT (Col-0) and the indicated mutants. Data represent means ± SD (*n* = 6-9 plants). Scale bars = 2 cm. **c-d** *ITPK1* expression (**c**) and shoot P_i_ levels (**d**) in 3-week-old WT (Col-0), *itpk1* and three independent *itpk1* lines transformed with *ITPK1* genomic DNA. Data represent means ± SD (*n* = 3 biological replicates in **c** and *n* = 8-9 plants in **d**). **e-h** Loss of ITPK1 results in P_i_ overaccumulation only in shoots but P_i_-independent root growth repression. Phenotypes (**e**), primary root length (**f**) and total P_i_ concentrations in shoots (**g**) or roots (**h**) of plants grown in agar plates with sufficient (625 μM P_i_) or deficient P_i_ supply (5 μM P_i_) for 7 days. Bars show means ± SD (*n* = 6 replicates with 3 plants each). Different letters indicate significant differences according to Tukey’s test (*P* < 0.05).

To investigate whether P_i_ accumulation is dependent on external P_i_ availability, we cultivated plants on agar plates supplemented with sufficient or insufficient P_i_. A root phenotypical analysis revealed that under sufficient P_i_ supply *itpk1* plants had shorter roots than WT plants, a phenotype that was also largely rescued in the recomplemented lines (Fig. 1e and 1f)^21^. The short-root phenotype of *itpk1* plants was also observed when plants were exposed to low P_i_ supply in agar plates (Fig. 1e and 1f) or cultivated in hydroponics (Suppl. Fig. 3), and was likely not associated with P_i_ overaccumulation, as the length of primary roots of *pho2-1* plants was similar to WT (Suppl. Fig. 3). Furthermore, we found that increased total P_i_ accumulation of *itpk1* plants was not detected in roots, while in shoots it was dependent on P_i_ availability (Fig. 1g and 1h). The shoot P_i_ overaccumulation phenotype of *itpk1* has been associated with defective down-regulation of PSRs in roots^13^. Our qPCR analysis confirmed that the expression of many PSI genes was up-regulated in *itpk1* roots but only when plants were grown under sufficient P_i_ or when P_i_-deficient plants were resupplied with P_i_ for 6 hours (Suppl. Fig. 4). Together, these results demonstrate that loss of ITPK1 but not ITPK2 affects the regulation of PSR in plants.

### ITPK1 is required for P_i_-dependent synthesis of PP-InsPs

Recently, two studies have raised strong evidence for the role of InsP_8_ in controlling P_i_ homeostasis in plants^14, 15^. To further investigate which InsPs respond to rapid changes in the plant’s P_i_ status, we first performed SAX-HPLC analyses of extracts from [^3^H]-inositol-labeled WT seedlings. Of all InsPs detected, only one InsP_3_ isomer, InsP_6_ and the PP-InsPs InsP_7_ and InsP_8_ decreased in response to P_i_ starvation and increased again when P_i_ was resupplied to P_i_-starved seedlings for 6 hours (Fig. 2a and 2b). Notably for InsP_8_, P_i_ resupply increased levels beyond those detected in seedlings continuously grown with sufficient P_i_ (Fig. 2b).

**Fig. 2.**
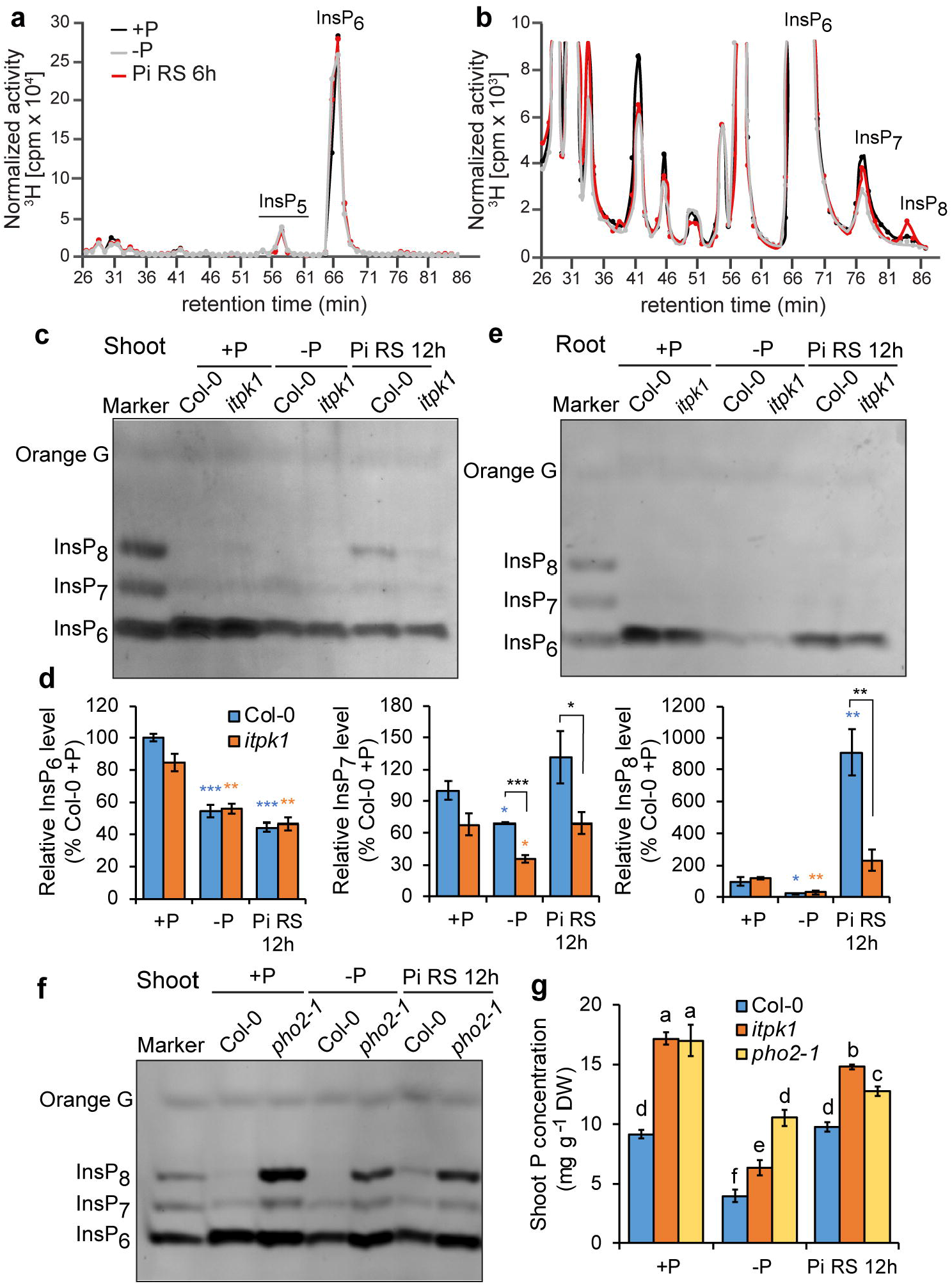
InsP_7_ and InsP_8_ levels in shoots respond to P_i_ availability in an ITPK1-dependent manner. **a-b** HPLC profiles of 17-day-old *Arabidopsis* WT (Col-0) seedlings radiolabeled with [^3^H]-*myo*-inositol. Seedlings were grown either with P_i_ (+P) or without P_i_ (-P) or resupplied with P_i_ for 6 h prior to harvest (Pi RS 6h). Full, normalized spectra (**a**) and zoom-in view of the same profile (**b**). The experiment was repeated two times with similar results, and representative results from one experiment are shown. **c-e** PAGE of InsP levels in shoots (**c**), quantification of PAGE signals from shoots (**d**) and PAGE of roots (**e**) of WT (Col-0) and *itpk1* plants. Plants were grown in hydroponics in P_i_-sufficient solution (+P), exposed for 4 days to P_i_ starvation (-P) or resupplied with P_i_ for 12 hours (Pi RS 12h). InsPs were eluted from TiO_2_ beads, separated by PAGE and visualized by Toluidine blue and DAPI. Data represent means ± SE (n = 3-4 biological replicates). \**P* < 0.05, \*\**P* < 0.01 and \*\*\**P* < 0.001 according to pairwise comparison with Student’s *t*-test. **f-g** InsP_7_ and InsP_8_ levels are strongly increased in the shoots of the P_i_-overaccumulating mutant *pho2-1*. PAGE of shoots (**f**) and shoot P_i_ levels in response to P_i_ deficiency and P_i_ resupply (**g**). Plants were cultivated as described in **c-e**. Data represent means ± SE (*n* = 4). Different letters indicate significant differences according to Tukey’s test (*P* < 0.05).

In order to increase the throughput of InsP_6_, InsP_7_ and InsP_8_ analysis and to allow assessing fully developed plants under more physiological conditions, we grew plants in hydroponics with aerated solution. Highly anionic InsPs and PP-InsPs extracted from plant tissues were purified by titanium dioxide (TiO_2_)-based pulldown followed by separation via PAGE and subsequent visualization by Toluidine Blue and 4′,6-diamidino-2-phenylindole (DAPI) based on previously established protocols^23, 24, 25^. In agreement with our HPLC analyses, InsP_6_, InsP_7_ and InsP_8_ signals in shoots of WT plants were strongly decreased when plants were deprived of P_i_ for 4 days (Fig. 2c and 2d). However, only InsP_7_ and especially InsP_8_ level were quickly restored by refeeding plants with P_i_ for 12 hours. Compared to WT, InsP_7_ and InsP_8_ levels were significantly lower in shoots of *itpk1* plants especially after P_i_ resupply (Fig. 2c and 2d). These results confirmed that ITPK1 functions as a cellular InsP_6_ kinase *in planta* and suggested that an ITPK1-dependent InsP_7_ pool is required and rate limiting for the efficient synthesis of InsP_8_ under conditions of high P_i_ availability. In contrast to *itpk1*, PP-InsP levels were not compromised in the *itpk2-2* mutant (Suppl. Fig. 5a), further suggesting that ITPK1 can compensate for the loss of *ITPK2*. Notably, in roots of both WT and *itpk1* plants, InsP_7_ and InsP_8_ were barely detected by PAGE (Fig. 2e).

To further address the role of different InsPs in PSR, we analyzed the *itpk4-1* mutant, which was recently reported to display reduced levels of InsP_5_[1/3-OH], InsP_6_ and InsP_7_^13^. Whereas HPLC analysis of [^3^H]-inositol-labeled seedlings confirmed the reported defects in InsP_5_ [1/-OH] synthesis^21^, our PAGE analyses did not detect any changes in InsP_6,_ InsP_7_ nor InsP_8_ levels in this mutant as compared to WT irrespective of the imposed P_i_ regime (Suppl. Fig. 5b and 5c). Thus, the undisturbed synthesis of PP-InsPs in *itpk4-1* plants may explain why this mutant shows no PSR phenotypes (Fig. 1b)^13^.

Considering that only InsP_7_ and InsP_8_ responded to P_i_, we then investigated the accumulation of these PP-InsPs in the P_i_ overaccumulator mutant *pho2-1*. We found that, compared to WT, *pho2-1* plants accumulated much higher levels of InsP_7_ and InsP_8_, even when grown on low P_i_ for 4 days (Fig. 2f). Notably, elemental analysis revealed that P_i_ levels in shoots of *pho2-1* plants exposed to low P_i_ were still significantly higher than those detected in WT plants grown under sufficient P_i_ (Fig. 2g). Together, these results indicate that the synthesis of PP-InsPs is dependent on the internal, cellular P_i_ levels rather than external P_i_ availability.

### Synthesis of InsP_7_ and InsP_8_ relies on ITPK1 and VIH2 and on InsP6 compartmentalization

As suggested in previous studies, the presence of InsP_7_ and InsP_8_ is likely central for P_i_ sensing in cells, as they promote the interaction of PHR with SPX proteins^10, 14, 15^. Intriguingly, our elemental analysis revealed that total P_i_ levels in shoots were not altered to the same degree in *vih2-4*, although previous HPLC analyses of [^3^H]-inositol-labeled seedlings indicated that InsP_8_ levels are strongly decreased in this mutant^17^. We further confirmed by PAGE the strong decrease of InsP_8_ in shoots of P_i_-resupplied *vih2-4* plants (Fig. 3a and 3b). PAGE analysis also detected a significant increase in InsP_7_ levels in *vih2-4* shoots after P_i_ resupply (Fig. 3a and 3b), suggesting that a certain InsP_7_ pool is regulated by VIH2-dependent conversion to InsP_8_ and that an increase of this pool in *vih2* mutant plants appears to partially compensate for the loss of InsP_8_ in regulating P_i_ signaling.

**Fig. 3.**
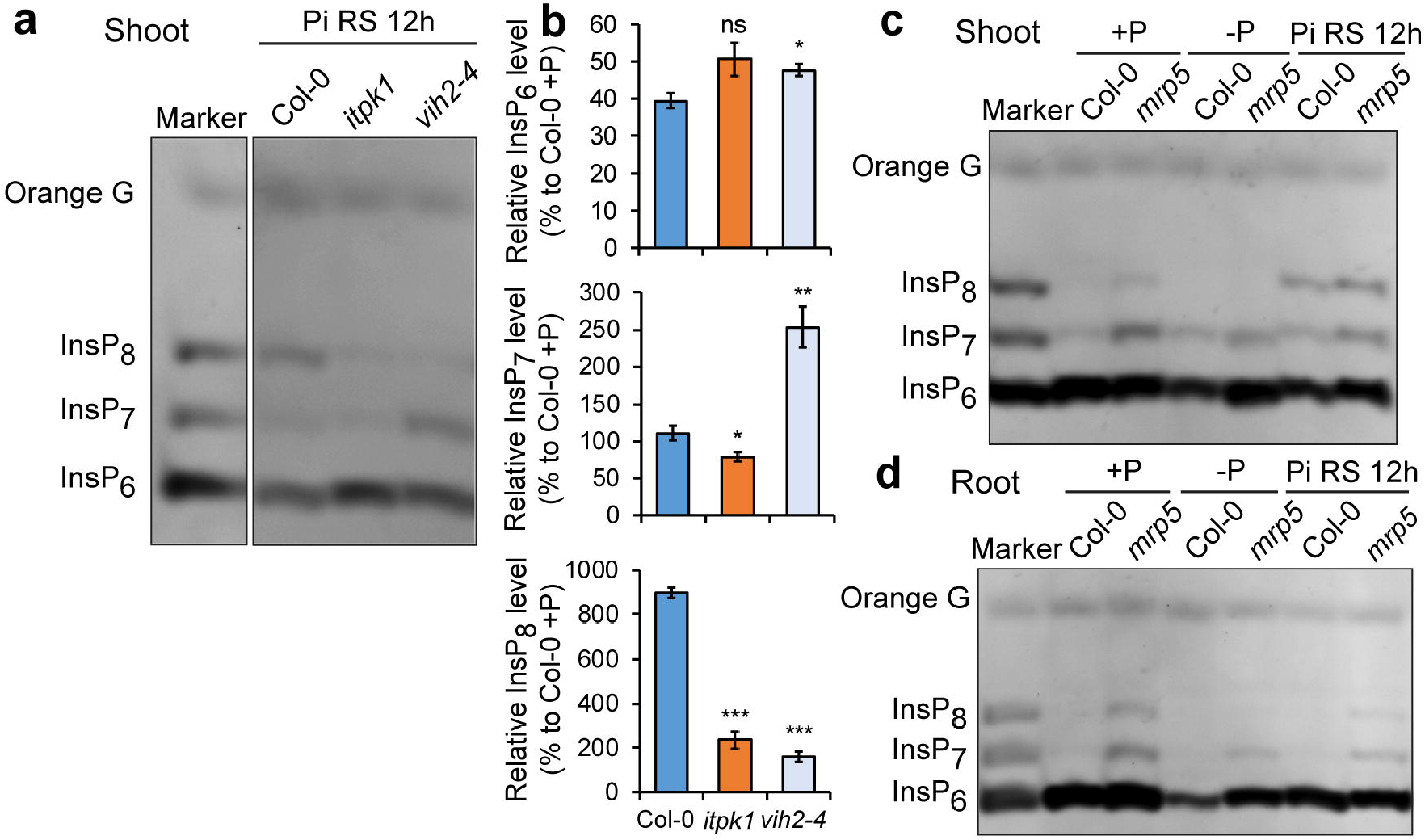
ITPK1-and VIH2-dependent synthesis of PP-InsPs and dependency on InsP_6_ compartmentation. **a-b** Loss of VIH2 increases InsP_7_ levels in shoots. InsP detection in shoots of *Arabidopsis* WT, *itpk1* and *vih2-4* plants (**a**) and quantification of PAGE signals (**b**) 12 h after P_i_ resupply to P_i_-starved plants in hydroponics. Data represent means ± SE (*n* = 4 biological replicates). \**P* < 0.05, \*\**P* < 0.01 and \*\*\**P* < 0.001 according to pairwise comparison with Student’s *t*-test. **c-d** Impaired InsP_6_ transport into the vacuole increases InsP_7_ and InsP_8_ levels in shoots and roots. InsP determination in shoots (**c**) and roots (**d**) of WT (Col-0) and *mrp5*. Plants were grown in hydroponics in P_i_ - sufficient solution (+P), exposed for 4 days to P_i_ starvation (-P) or resupplied with P_i_ for 12 hours (Pi RS 12h).

*ITPK1, VIH1* and *VIH2* are not only expressed in shoots but also in roots^13, 15^. However, since little to no InsP_7_ and InsP_8_ could be detected by PAGE in roots of Arabidopsis and rice plants (Fig. 2e and Suppl. Fig. 5), we wondered whether the availability of cytosolic InsP_6_ could determine the amount of InsP_7_ and InsP_8_ that can be synthesized. To test this hypothesis, we assessed these PP-InsPs in shoots and roots of *mrp5*, a mutant defective in vacuolar loading of InsP_6_^26^. In contrast to WT, we could detect InsP_7_ and InsP_8_ both in shoots and in roots of *mrp5* plants (Fig. 3c and 3d). Lack of *MRP5* resulted in higher levels of InsP_7_ and InsP_8_ as compared to WT but InsP_8_ was still clearly responsive to P_i_ starvation and P_i_ resupply in this mutant. However, increased levels of these PP-InsPs in roots did not significantly affect P_i_ accumulation in *mrp5* shoots (Fig. 1b). Altogether, these results demonstrate that P_i_-dependent synthesis of InsP_7_ and InsP_8_ relies on stepwise phosphorylation reactions mediated by ITPK1 and VIHs, and on the cytosolic/nucleoplasmic availability of InsP_6_.

### ITPK1 has InsP_6_ kinase and ATP synthase activities

Recently, we demonstrated that recombinant ITPK1 phosphorylates InsP_6_ *in vitro* at position 5 thereby generating 5-InsP_7_, which is the main InsP_7_ isomer detected in seeds^20^. Considering that ITPK1 is required for the robust increase in InsP_8_ when P_i_-deficient plants are resupplied with P_i_ (Fig. 2c and 2d), we asked whether ITPK1 is able to function as an InsP_7_ kinase. As shown in Suppl. Fig. 7a, neither of the natural occurring isomers, 1-InsP_7_ and 5-InsP_7_, appears to be a substrate for ITPK1 kinase activity suggesting that the ITPK1-dependent increase in InsP_8_ after P_i_ resupply is caused by an increase in a rate limiting pool of InsP_7_. To further investigate the enzymatic properties of ITPK1, we performed nuclear magnetic resonance (NMR) assays with the recombinant protein taking advantage of [^13^C_6_]-labelled InsP_6_. First, InsP_6_ kinase reaction conditions were analyzed with respect to magnesium ion (Mg^2+^) concentration and temperature dependencies as well as to quenching efficiency by EDTA (Suppl. Fig. 7b-d). Using 2.5 mM ATP, optimal enzyme activity was observed with as little as 2.5 mM Mg^2+^ and conversion could be fully quenched by a surplus of EDTA exceeding the Mg^2+^ concentration by 1.36 mM. Even though the protein behaved well at 37°C and increased kinase activities were observed at this elevated temperature (Suppl. Fig. 7d), we decided to carry out subsequent experiments at 25°C to more closely reflect temperatures in unstressed plants. Subsequent kinetic analysis revealed that ITPK1 exhibits a surprisingly high K_M_ for ATP of approximately 520 μM (Fig. 4a and 4b). Unlike VIHs, the kinase activity of ITPK1 was largely insensitive to P_i_ and was also not affected by the non-metabolizable P_i_ analog phosphite (Suppl. Fig. 8). When InsP_5_ [2-OH] was presented as substrate to ITPK1, no conversion could be detected (Suppl. Fig. 7e), suggesting that ITPK1 has no IPK-like activity to generate InsP_6_ from InsP_5_ [2-OH]. Furthermore, in contrast to InsP_6_ kinases of the IP6K/Kcs1 family, no activity was observed when 1-InsP_7_ was used as a substrate (Suppl. Fig. 7f and 7g), thus confirming our PAGE analysis (Suppl. Fig. 7a).

**Fig. 4.**
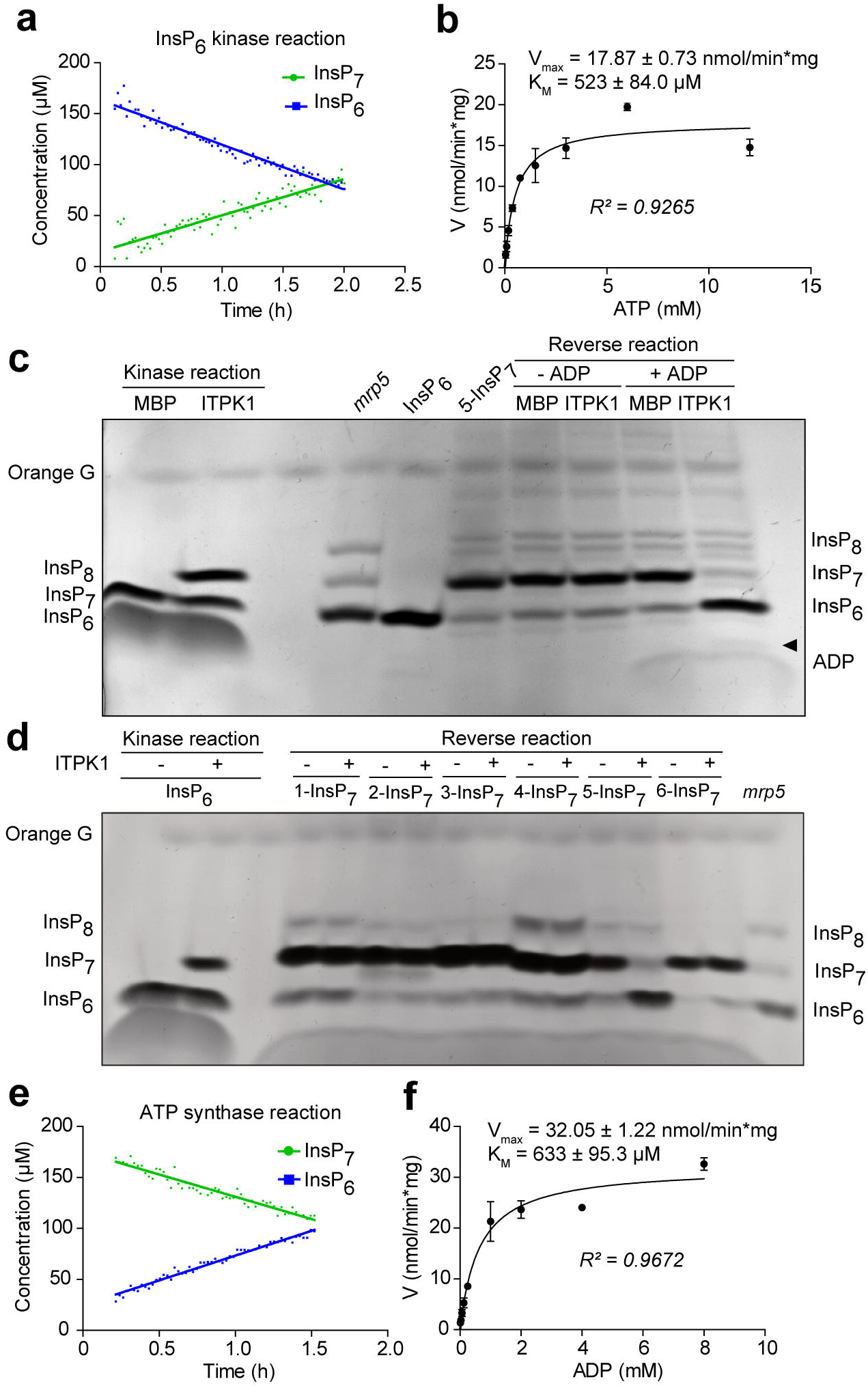
*In vitro* characterization of ITPK1 activity. **a-b** NMR analysis of InsP6 kinase activity of recombinant *Arabidopsis* ITPK1. Time-dependent conversion of InsP_6_ to 5-InsP_7_ (**a**) and reaction velocity determined at varying ATP concentrations (**b**). K_M_ and V_max_ were obtained after fitting of the data against the Michaelis-Menten model. **c-d** In the presence of ADP, recombinant *Arabidopsis* ITPK1 mediates 5-InsP_7_ hydrolysis (**c**) but not the hydrolysis of other InsP_7_ isomers (**d**). InsPs were separated via PAGE and visualized by Toluidine Blue staining. The identity of bands was determined by migration compared to InsP_6_ and 5-InsP_7_ standards and TiO_2_-purified *mrp5* seed extract. InsP_6_ kinase reaction served as positive control for the reverse reactions. Purified His_8_-MBP tag (MBP) served as negative control for ITPK1. Arrowhead in **c**, indicates the presence of a small ATP band just above ADP. **e-f** NMR analysis of reverse reaction of recombinant *Arabidopsis* ITPK1. Accumulation of InsP_6_ and conversion of 5-InsP_7_ (**e**) and reaction velocity determined at varying ADP concentrations (**f**). K_M_ and V_max_ were obtained after fitting of the data against the Michaelis-Menten model.

The characterization of structurally and sequence-unrelated mammalian InsP_6_ kinases of the IP6K family has demonstrated that these enzymes can shift their activities from kinase to ADP phosphotransferase at low ATP-to-ADP ratios^27, 28^. This prompted us to assess if also ITPK1 possesses such activity. *In vitro* reactions with unlabeled 5-InsP_7_ and subsequent PAGE analyses revealed that ITPK1 indeed mediates 5-InsP_7_ hydrolysis and that this activity only occurs in the presence of ADP (Fig. 4c). A similar activity using any other InsP_7_ isomer as phosphoryl donor could not be detected (Fig. 4d), suggesting a high degree of substrate specificity not only for ITPK1-mediated kinase activity but also for the reverse reaction (i.e., InsP_7_ dephosphorylation / ATP synthesis). To determine the kinetic parameters of this reaction, we subsequently incubated ITPK1 with [^13^C_6_]-labelled 5-InsP_7_ in the presence of ADP and detected the formation of ATP and InsP_6_ (Suppl. Fig. 9a and 9b). No ATP formation was detected when the enzyme was incubated without 5-InsP_7_ (Suppl. Fig. 10c). Interestingly, the velocity of the reverse reaction was almost two times faster than the forward reaction, whereas the K_M_ for ADP and ATP was relatively similar for ITPK1 (Fig. 4b, 4e and 4f). Taken together, these results demonstrate that ITPK1-mediated InsP_6_ kinase and ADP phosphotransferase activities are regulated by adenylate energy charge. In agreement with results obtained in agar plate-grown seedlings^15^, we observed that ATP levels and ATP/ADP ratios dropped significantly in response to P_i_ deficiency in shoots of hydroponically-grown WT plants but rapidly increased after P_i_ resupply (Suppl. Fig. 10). Thus, the P_i_-dependent changes in energy status of plants may ultimately regulate the synthesis of InsP_7_ by shifting the activity of ITPK1.

### ITPK1 activity in shoots controls PHR1- and PHL1-regulated PSRs

To directly address whether ITPK1-dependent synthesis of PP-InsPs in shoots or roots is responsible for P_i_ accumulation, we performed grafting experiments. As expected, shoot P_i_ overaccumulation was also detected when roots and shoots of *itpk1* plants were self-grafted (Fig. 5a). However, shoot P_i_ was almost back to WT levels when Col-0 shoots were grafted onto *itpk1* roots, while remaining approximately 75% higher when *itpk1* shoots were grafted onto Col-0 roots (Fig. 5a). Shoot dry weight or shoot levels of other nutrients were not significantly altered by the different graft combinations (Suppl. Fig. 11). These results suggest that ITPK1 activity in shoots is more determinant for the regulation of PSR and P_i_ accumulation.

**Fig. 5.**
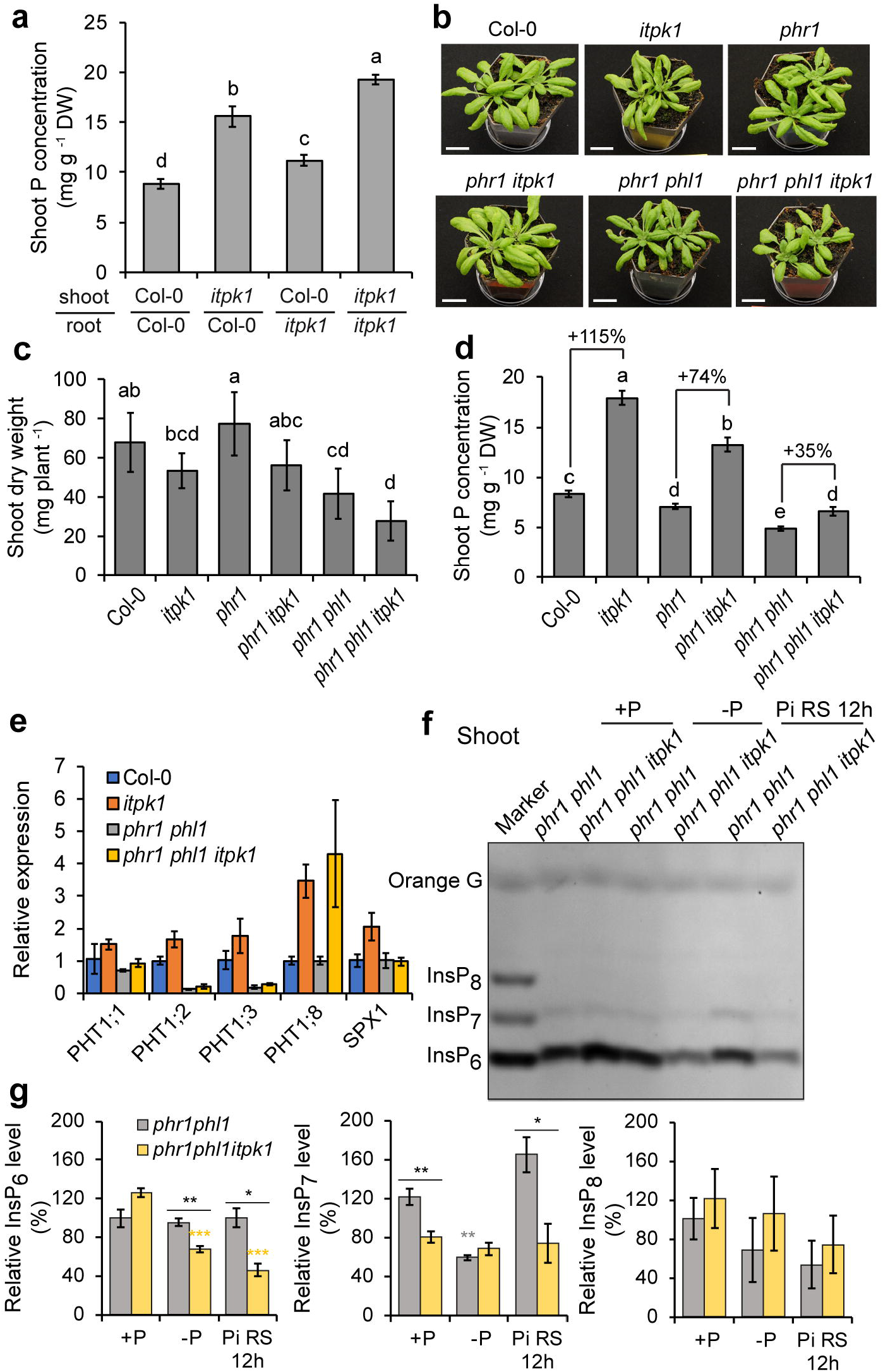
P_i_ starvation responses are mainly regulated by ITPK1 activity in shoots. **a** Total P_i_ concentration in shoots of self-grafted or reciprocally grafted WT (Col-0) and *itpk1*. Plants were grafted on agar plates and left recovering for 2 weeks. Positive grafts were transferred to peat-based substrate for another 2 weeks. Data represent means ± SD (*n* = 5-7 plants). **b-d** Genetic interplay between PHR1/PHL1 and ITPK1 in P_i_ sensing. Phenotype (**b**), shoot dry weight (**c**) and shoot P_i_ levels (**d**) of 3-week-old WT and indicated mutants grown on peat-based substrate. Data represent means ± SD (*n* = 6 plants). In **a**, **c** and **d**, letters indicate significant differences according to Tukey’s test (*P* < 0.05). **e** ITPK1-dependent expression of P_i_ deficiency-induced genes in roots of the indicated P_i_-sufficient plants. Data represent means ± SE (*n* = 4 replicates). **f-g** PHR1- and PHL1-dependent synthesis of InsPs as revealed by PAGE (**f**) and relative quantification of signals (**g**) in indicated double and triple mutants grown in hydroponics with P_i_-sufficient solution (+P), exposed to 4 days of P_i_ starvation (-P) or resupplied with P_i_ for 12 hours (Pi RS 12h). Data represent means ± SE (*n* = 4 biological replicates). \**P* < 0.05, \*\**P* < 0.01 and \*\*\**P* < 0.001 according to pairwise comparison with Student’s *t*-test.

We then addressed the putative involvement of ITPK1 in P_i_ signaling by analyzing the genetic interaction between ITPK1 and the transcription factors PHR1 and PHL1, which together control most of the transcriptional responses induced by P_i_ starvation^2^. Shoot growth was not significantly altered in homozygous double or triple mutants as compared to single or double mutants (Fig. 5b and 5c). Although *phr1 itpk1* and *phr1 phl1 itpk1* plants still accumulated significantly more P_i_ than *phr1* and *phr1 phl1*, respectively, the relative increments were smaller than in the presence of functional PHR1 and PHL1 (Fig. 5d). In turn, the short-root phenotype caused by *ITPK1* mutation could not be restored by knocking out these transcription factors (Suppl. Fig. 12). While most P_i_ starvation-induced transcriptional responses were also suppressed in the triple mutant, absence of ITPK1 maintained *PHT1;8* up-regulated in the *phr1 phl1* background (Fig. 5e) potentially suggesting a PHR1/PHL1-independent regulation of *PHT1;8* expression. Finally, PAGE analysis revealed that InsP_8_ levels remained very low when these two transcriptional regulators were knocked out (Fig. 5f and 5g). In contrast, P_i_ resupply-induced InsP_7_ accumulation was largely independent of PHR1 and PHL1. Collectively, these observations demonstrate that ITPK1-dependent PP-InsP synthesis in shoots is required for undisturbed coordination of systemic P_i_ signaling by PHR1 and PHL1.

### P_i_-dependent synthesis of PP-InsPs is conserved across species

Next, we assessed whether P_i_-dependent regulation of InsP_7_ and InsP_8_ is evolutionarily conserved across species. Similar to Arabidopsis, we observed that both InsP_7_ and InsP_8_ levels decreased strongly in response to P_i_-deficiency in shoots of hydroponically-grown rice plants and were quickly restored by refeeding plants with P_i_ (Fig. 6a and 6b). Notably, in this species, a clear InsP_7_ signal could be observed as soon as 30 min after P_i_ resupply and appeared before substantial changes in InsP_8_ could be detected by PAGE. We also detected increased accumulation of PP-InsPs in P_i_-replete and P_i_-refed gametophores of the moss *Physcomitrella patens* (Fig. 6c-6e), suggesting that P_i_-dependent InsP_7_ and InsP_8_ synthesis is also conserved in non-vascular land plants.

**Fig. 6.**
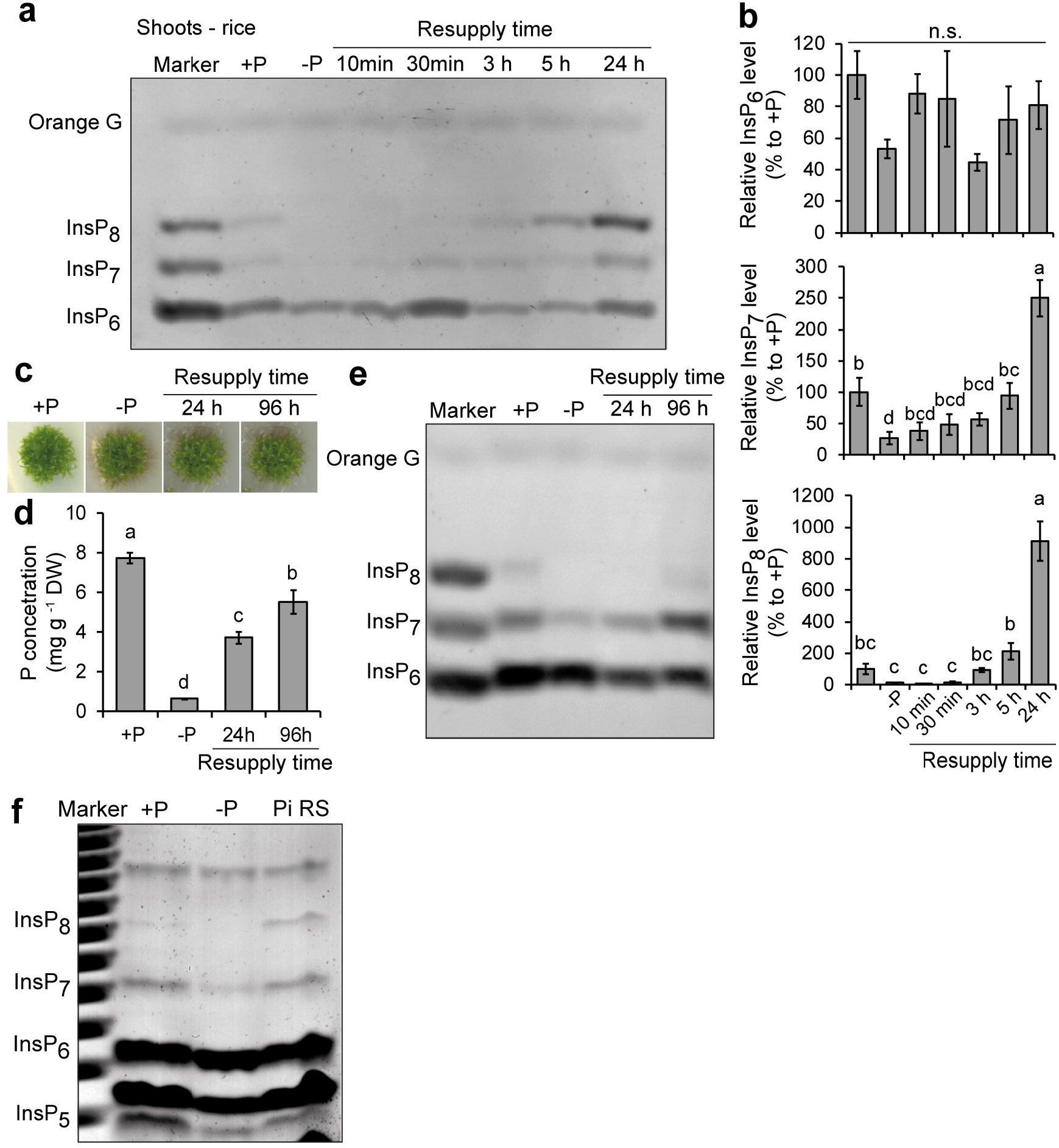
P_i_-dependent regulation of InsP_8_ levels is conserved in multicellular organisms. **a-b** Time-course analysis of InsPs in response to P_i_ starvation and P_i_ resupply in rice shoots. PAGE (**a**) and relative quantification of bands (**b**) of rice plants cv. Nipponbare grown in hydroponics. Data represent means ± SE (*n* = 4 biological replicates). Different letters indicate significant differences according to Tukey’s test (*P* < 0.05). **c-e** Phenotype (**c**), total P_i_ levels (**d**) and PAGE of P_i_-dependent synthesis of InsP_7_ and InsP_8_ (**e**) in gametophores of *Physcomitrella patens*. Plants were cultivated on sufficient P_i_ (+P), starved of P_i_ for 30 days (-P) or resupplied with P_i_ for the indicated time. Data represent means ± SD (*n* = 3 biological replicates). Different letters indicate significant differences according to Tukey’s test (*P* < 0.05). **f** PAGE analysis of wild-type HCT116 cell extracts during P_i_ starvation and resupply. Cells were starved in P_i_-free media for 18 h and resupplied with P_i_ for 3.5 h. Cells were harvested at the same time. The experiment was repeated twice with similar results.

Since we recently showed that purified recombinant human ITPK1 is also able to catalyze 5-InsP_7_ synthesis^20^, we assessed P_i_-dependent synthesis of PP-InsPs in the human HCT116 cell line. We found that while InsP_6_ levels remained largely unaffected by P_i_ conditions, both InsP_7_ and InsP_8_ decreased in cells after removing P_i_ from the culture and sharply increased again after P_i_ resupply (Fig. 6f). Altogether, these results indicate that P_i_-dependent synthesis of InsP_7_ and InsP_8_ seems to be evolutionary conserved across a range of different multicellular species.

## DISCUSSION

Previously, isothermal titration calorimetry experiments demonstrated that 5-InsP_7_ interacted more strongly with SPX domains than P_i_ and that 5-InsP_7_ concentrations as low as 20 μM can effectively promote the interaction of OsSPX4 with OsPHR2^10^, suggesting that InsP_7_ can affect P_i_ signaling. The finding that an *itpk1* mutant in Arabidopsis over-accumulates P_i_ and constitutively express P_i_ starvation-induced genes under sufficient P_i_ (Fig. 1b; Suppl. Figs. 2 and 4)^13^ provided further evidence that ITPK1 function is required for proper regulation of P_i_ homeostasis. In this work, we demonstrate that InsP_7_ and InsP_8_ most closely mirror P_i_ levels in different organisms as they decrease in response to P_i_ deficiency and quickly increase after P_i_ resupply (Fig. 2a-2c; Fig. 6). In *A. thaliana*, we show that both InsP_7_ and InsP_8_ levels are compromised in shoots of *itpk1* plants, especially when P_i_-deficient plants are resupplied with P_i_ (Fig. 2c and 2d). Since we detected only InsP_6_ kinase but no InsP_7_ kinase activity with purified recombinant ITPK1 (Fig. 4c; Suppl. Fig. 7a)^20^, the decreased InsP_8_ levels in *itpk1* plants likely resulted from the diminished 5-InsP_7_ available for the subsequent phosphorylation at the C1 position by VIH1 and VIH2. Our results also indicate that ITPK1 can shift its activity and become an ADP phosphotransferase that specifically hydrolyses 5-InsP_7_ when adenylate energy charges are low, such as in P_i_-deficient cells (Fig. 4; Suppl. Fig. 10). Thus, our results with ITPK1 and those of Zhu et al.^15^ with VIH1 and VIH2 provide a detailed view on how changes in cellular P_i_ status alter the activity of these enzymes to efficiently produce and eliminate the signaling molecules InsP_7_ and InsP_8_ in plants (Fig. 7).

**Fig. 7.**
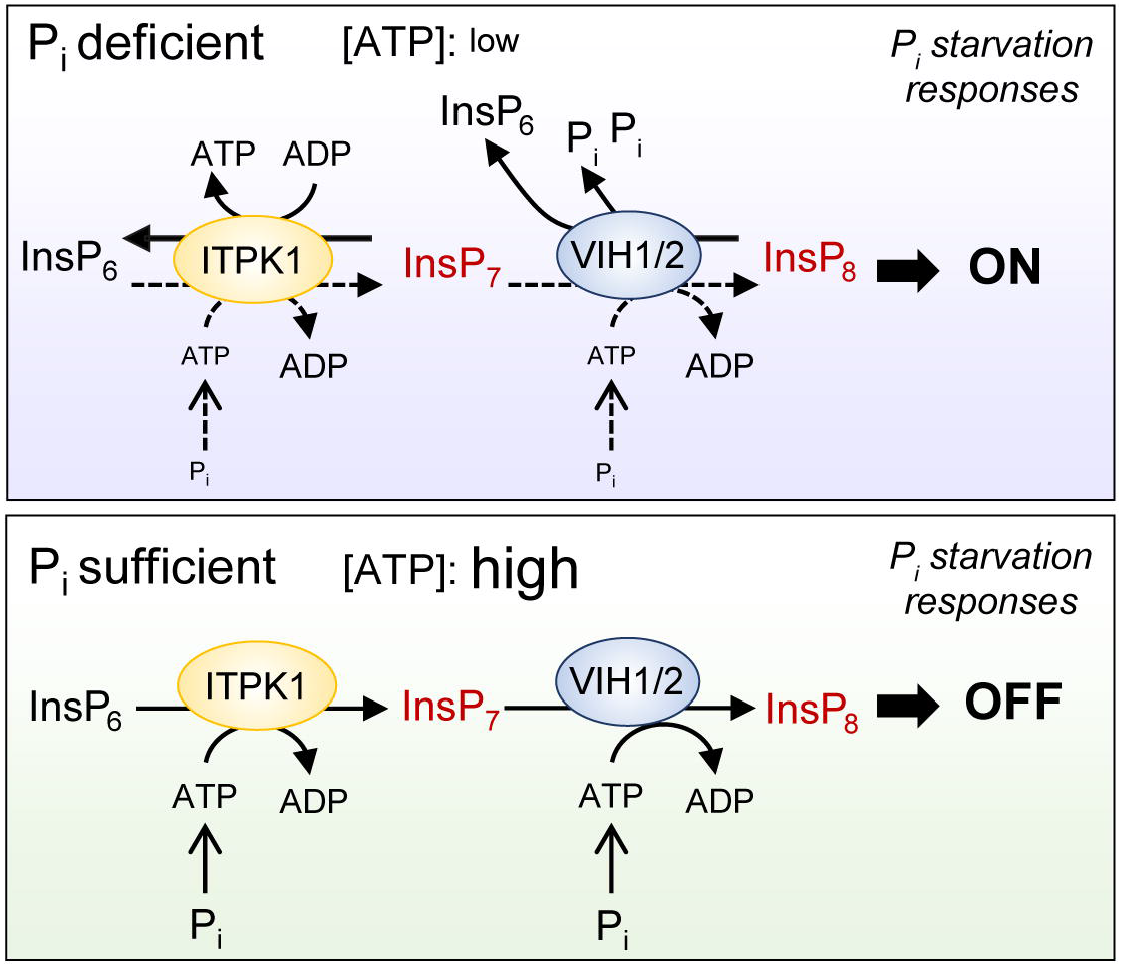
Model for ITPK1-dependent generation and removal of InsP_7_ and its link with VIHs and P_i_ signaling. In P_i_-deficient cells, low ATP levels stimulate ITPK1 to catalyze P_i_ transfer from InsP_7_ to ADP, thereby generating ATP and decreasing InsP_7_. Decreased ATP and P_i_ levels also activate the pyrophosphatase activity of VIHs to break down InsP_8_. The removal of PP-InsPs destabilizes the association between PHRs and SPXs, allowing PHRs to induce P_i_ starvation responses. When cells regain sufficient P_i_, which increase ATP levels, ITPK1-mediated InsP_6_ kinase activity is stimulated and the reverse reaction towards InsP_7_ is inhibited. InsP_7_ generated by ITPK1 serves then as substrate for InsP_8_ production via the kinase domain of VIHs. As a consequence of increased PP-InsPs, SPX proteins recruit PHRs to repress P_i_ starvation responses.

Although both ITPK1 and ITPK2 possess InsP_6_ kinase activity^20^, PSR seems only disturbed in *itpk1* but not *itpk2* mutants (Fig. 1b)^13^. PP-InsPs are not significantly altered in *itpk2* seedlings (Suppl. Fig. 5a)^21^, suggesting that ITPK1 can fully compensate for the loss of ITPK2. Furthermore, since only *ITPK2* transcription is induced by P_i_ deficiency (Suppl. Fig. 13), it is likely that ITPK1 and ITPK2 play independent roles in P_i_ signaling. In a previous study, the putative role of InsP_7_ in plant P_i_ signaling was excluded because HPLC analysis of [^32^P]-P_i_-labeled plants showed that InsP_7_ levels were similarly compromised in *itpk1* and *itpk4* plants^13^, whereas only *itpk1* displays obvious PSR defects (Fig. 1b)^13^. However, our PAGE analyses showed that InsP_7_ and InsP_8_ levels are not compromised in the *itpk4-1* mutant (Suppl. Fig. 5b and 5c). Therefore, a role for InsP_7_ and/or InsP_8_ cannot be excluded based on the comparison of *itpk1* and *itpk4-1*. Meanwhile, two recent studies raised compelling evidence that InsP_8_ acts as a signaling molecule to regulate P_i_ homeostasis in Arabidopsis, as *vih1 vih2* double mutants exhibit constitutive up-regulation of PSI genes and strong P_i_ hyperaccumulation^14, 15^. Here, we show that a P_i_-dependent accumulation of InsP_8_ and InsP_7_ appears to be conserved in land plants, including dicotyledonous and monocotyledonous vascular species as well as non-vascular species (Fig. 2a-2c; Fig. 6a-e). Of note, our PAGE results from independent experiments demonstrate that InsP_8_ is more sensitive to P_i_ than InsP_7_. However, InsP_7_ levels were clearly reduced when plants were exposed to more prolonged periods of P_i_ starvation (Fig. 6a, 6b and 6e). Furthermore, we also found that not only InsP_7_ but also InsP_8_ is induced by P_i_ in human HCT116 cells (Fig. 6f). P_i_-dependent regulation of InsP_8_ in human cells has likely also an impact on P_i_ homeostasis, as this PP-InsP was recently shown to be functionally dominant over 5-InsP_7_ and 1-InsP_7_ in the regulation of the cellular P_i_ exporter protein Xenotropic and Polytropic Retrovirus Receptor 1 (XPR1)^29^.

In Arabidopsis, InsP_8_ levels are strongly reduced while InsP_7_ significantly increased in the *vih1 vih2* double mutant^15^, suggesting that InsP_7_ itself plays only a relatively minor role in P_i_ signaling. However, considering the evidence for the involvement of InsP_8_ in other cellular processes^17, 30, 31^, it might well be that the severe growth defects of *vih1 vih2* plants^14, 15^ are not solely related to disturbed P_i_ homeostasis. Here, we analyzed the *vih2-4* single mutant, which grows similarly to the WT (Fig. 1a), and found that whereas InsP_8_ levels in *vih2-4* plants were as low as in *itpk1*, strong P_i_ overaccumulation was only observed for *itpk1* plants (Fig. 1b; Fig. 3a and 3b). Since InsP_8_ reduction in *vih2-4* plants is accompanied by increased InsP_7_ levels (Fig. 3a and 3b), it is likely that, under certain circumstances, InsP_7_ may partially compensate for the loss of InsP_8_. Noteworthy, also InsP_7_ is involved in other cellular processes, such as auxin perception, as 5-InsP_7_ can bind to the auxin receptor complex^21^. Importantly, the role of ITPK1 in auxin perception is largely independent of its function in P_i_ homeostasis^21^, suggesting that different tissue-specific InsP_7_ pools may regulate diverse signaling pathways. This assumption is supported by the fact that the auxin-associated short-root phenotype of *itpk1* plants cannot be complemented with P_i_ nor attenuated by disrupting PHR1 and PHL1 (Fig. 1e and 1f; Suppl. Figs. 3 and 12). However, to directly assess the contribution of InsP_7_ to different physiological processes, it would be necessary to develop a strategy to specifically disrupt InsP_7_ synthesis without altering InsP_8_ levels.

The genetic analysis of *phr1 itpk1* and *phr1 phl1 itpk1* multiple mutants demonstrated that, in the absence of PHRs, *ITPK1* mutation did not anymore mis-regulate the expression of P_i_ starvation-induced genes nor result in uncontrolled P_i_ accumulation under sufficient P_i_ (Fig. 5d and 5e). These results place ITPK1 upstream of PHRs and further support that P_i_-dependent synthesis of InsP_7_ by ITPK1 and of InsP_8_ by VIHs is critical for undisturbed P_i_ signaling in plants. Our grafting experiment further demonstrated that P_i_ overaccumulation in *itpk1* was mainly due to missing ITPK1 activity in shoots rather than in roots (Fig. 5a), thus pointing to a major role of PP-InsPs in P_i_ signaling in shoots. This result was somewhat surprising, since *ITPK1*, *VIH1* and *VIH2* are expressed in shoots and roots^13, 15, 17^. However, in line with earlier indications from [^32^P]-P_i_-labeling^13^, our PAGE analyses also detected much higher InsP_7_ and InsP_8_ levels in shoots than in roots of both Arabidopsis and rice plants (Fig. 2c and 2e; Suppl. Fig. 5). However, accumulation of these PP-InsPs in roots could be induced by disturbing MRP5-mediated InsP_6_ loading into the vacuole (Fig. 3d). Nonetheless, it remains unknown whether InsP_6_ availability to ITPK1 in leaves is also regulated by P_i_. Whereas ITPK1 is located both in the nucleus and in the cytoplasm^13^, VIH1 and VIH2 are reported to display cytoplasmic localizations^15^. Given that SPXs act as receptors for PP-InsPs^10, 14, 32, 33, 34^, ligand availability controls the interaction between SPXs and PHRs. To affect P_i_ signaling, PP-InsPs most likely have to cross the nuclear envelope, especially because SPX1, SPX2 and SPX3 are localized in the nucleus^4, 35^. Alternatively, they may also interact with proteins located in the cytoplasm, such as SPX4^36^.

The emerging model is that in the presence of low levels of PP-InsPs ligands, as when plants suffer from P_i_ deficiency (Fig. 2a-2d; Fig 6a-6e), SPX receptors are unable to bind to PHRs. Consequently, these transcription factors are free to bind to the P1BS motifs present in the promoters of several P_i_-responsive genes^1, 2^. Our findings that InsP_7_ and InsP_8_ levels quickly increase after P_i_ resupply in different plant species (Fig. 2a-2d; Fig. 6a-6e) provide insights into how this mechanism can be reversed once P_i_-starved plants regain access to P_i_. It is likely that the increased ligand levels shortly overlap with high abundance of SPX proteins, allowing for the formation of SPX - PHR complexes and the subsequent quick inhibition of PHR-dependent transcriptional responses.

Our PAGE analyses demonstrate that InsP_7_ and InsP_8_ levels dynamically reflect the P_i_ status in different organisms, indicating that PP-InsPs synthesis and degradation (or simply turnover) must be tightly controlled. P_i_-dependent accumulation of InsP_8_ has been proposed to rely on the bifunctional activity of the diphosphoinositol pentakisphosphate kinases VIH1 and VIH2^14, 15^. The relative kinase and phosphatase activities of this type of bifunctional enzymes can be shifted according to cellular ATP and P_i_ levels. Indeed, purified Vip1 from yeast exhibited kinase activity and, hence, increased InsP_8_ synthesis at higher ATP concentrations, whereas at low ATP levels the protein functioned mainly as an InsP_8_ phosphatase^15^. In this context, the phosphatase activity of diphosphoinositol pentakisphosphate kinases is further inhibited by P_i_ itself^15, 22^. However, unlike VIHs, ITPK1 does not harbor the bifunctional kinase domain - phosphatase domain architecture but only the atypical ‘ATP-grasp fold’ ATP-binding site. Here, we demonstrate with independent approaches that ITPK1-mediated InsP_6_ conversion to 5-InsP_7_ depends on ATP availability (Fig. 4). Our NMR-based kinetic assays also revealed that Arabidopsis ITPK1 has a high K_M_ of approximately 523 μM for ATP (Fig. 4b), which is similar to the K_M_ recorded for mammalian IP6Ks^27, 28^, suggesting that ITPK1-dependent InsP synthesis will be compromised at low cellular adenylate energy charge.

In metazoans and yeast, PP-InsPs act as energy sensors and metabolic messengers^37, 38^, and fluctuations in ATP levels have been shown to correlate with changes in intracellular concentrations of InsP_7_^39^. In our conditions, we observed that, compared to P_i_-deficient plants, ATP levels and ATP/ADP ratios are significantly higher in plants grown continuously with sufficient P_i_ or shortly after P_i_ is resupplied to P_i_-deficient plants (Suppl. Fig. 10). Previous reports have indeed shown that in plants, yeast and human cells ATP levels drop in response to P_i_ starvation^15, 22, 40, 41^. Interestingly, we found that ITPK1 has also a high K_M_ for ADP and, in the presence of this adenine nucleotide, converted 5-InsP_7_ to InsP_6_ generating ATP in the process (Fig. 4c, 4d and 4e; Suppl. Fig. 9a). This activity is reminiscent of the ATP synthase activity recorded for mammalian IP6K-type InsP_6_ kinases^27, 28^ and even for the human diphosphoinositol pentakisphosphate kinase PPIP5K2^42^. Our kinetic analyses demonstrated that the enzyme has comparable K_M_ values for ATP and ADP and similar V_max_ values, i.e. similar efficiencies for the forward and reverse reactions (Fig. 4b and 4f), suggesting that relative adenylate energy charge determines whether ITPK1 phosphorylates InsP_6_ or synthesizes ATP from 5-InsP_7_. Consequently, ITPK1 activity would allow cells to, for example, rapidly remove the signaling molecule InsP_7_ in low P_i_ conditions, when ATP levels and ATP/ADP ratios decrease. This type of deactivation of InsP_7_/InsP_8_ signaling could bypass the requirement for dedicated PP-InsP hydrolases which are likely to slow down quick dynamic changes in PP-InsPs to induce jasmonate-related responses during wound response or insect attack^17^, or when P_i_ becomes suddenly available (Fig. 2a-2d; Fig. 6). While activation of the reverse reaction is unlikely to significantly alter global cellular ATP pools, localized ATP synthesis could quickly suppress InsP_7_-mediated P_i_ signaling and, at the same time, might buffer the adenylate energy charge in the vicinity of ITPK1 under conditions of low energy or low P_i_ supply. Thus, adenylate charge-driven changes in the activities of ITPK1 and VIHs may represent one underlying mechanism by which the cellular P_i_ status is transduced into specific PP-InsPs levels to regulate downstream signaling events.

## METHODS

### Arabidopsis thaliana: plant materials and growth conditions

Seeds of *Arabidopsis thaliana* T-DNA insertion lines *itpk1* (SAIL_65_D03), *itpk2-2* (SAIL_1182_E03), *itpk3* (SALK_120653), *itpk4-1* (SAIL_33_G08), *ipk1-1* (SALK_065337C), *vih2-4* (GK-080A07), *mips1* (SALK_023626), *mips2* (SALK_031685), *mips3* (SALK_120131), *mrp5* (GK-068B10), *pho2-1* (EMS mutant described previously^43^) and *phr1* (SALK_067629) were obtained from The European Arabidopsis Stock Centre (http://arabidopsis.info/). The *phr1 phl1* double mutant used in this study was described previously^12^. To generate the *phr1 itpk1* double and the *phr1 phl1 itpk1* triple mutant, we crossed *itpk1* (−/−) with, respectively, *phr1* (−/−) and the homozygous *phr1 phl1* mutant. F2 and F3 plants were genotyped by PCR using the primers indicated in Suppl. Table 1 to identify homozygous lines. Transgenic lines expressing the genomic *ITPK1* fragment in the *itpk1* background were generated as described previously^21^.

For sterile culture, Arabidopsis seeds were surface sterilized in 70% (v/v) ethanol and 0.05% (v/v) Triton X-100 for 15 min and washed twice with 96% (v/v) ethanol. Sterilized seeds were sown onto modified half-strength Murashige and Skoog (MS) medium^44^ containing 0.5% sucrose, 1 mM MES pH 5.5 and solidified with 1% (w/v) Difco agar (Becton Dickinson). After 7 days of preculture, seedlings were transferred to vertical plates containing fresh solid media supplemented with either sufficient P_i_ (625 μM P_i_) or low P_i_ (5 μM P_i_). Plants were grown under axenic conditions in a growth cabinet under the following regime: 10/14 h light/dark; light intensity 120 μmol m^−2^ s^−1^ (fluorescent lamps); temperature 22°C/18°C.

For hydroponic culture, Arabidopsis seeds were pre-cultured on rock wool moistened with tap water. After 1 week, tap water was replaced by half-strength nutrient solution containing 2 mM NH_4_NO_3_, 1 mM KH_2_PO_4_, 1 mM MgSO_4_, 1 mM KCl, 250 μM K_2_SO_4_, 250 μM CaCl_2_, 100 μM Na-Fe-EDTA, 30 μM H_3_BO_3_, 5 μM MnSO_4_, 1 μM ZnSO_4_, 1 μM CuSO_4_ and 0.7 μM NaMoO_4_ (pH adjusted to 5.8 by KOH). After 7 days, nutrient solution was changed to full-strength and replaced once a week (first 3 weeks), twice a week in the fourth week, and every 2 days in the following weeks. Aeration was provided to roots from the third week onwards. To induce P_i_ deficiency, KH_2_PO_4_ was replaced by KCl and P_i_ resupply was performed by refeeding P_i_-starved plants with 1 mM KH_2_PO_4_ for 12 h. Plants were grown hydroponically in a growth chamber under the above-mentioned conditions except that the light intensity was 200 μmol photons m^−2^ s^−1^ and supplied by halogen lamps.

Phenotypic characterization in soil substrate was performed by germinating seeds directly in pots filled with peat-based substrate (Klasmann-Deilmann GmbH, Germany). The pots were placed inside a conditioned growth chamber with a 22°C/18°C and 16-h/8-h light/dark regime at a light intensity of 120 μmol photons m^−2^ s^−1^ supplied by fluorescent lamps. Plants were bottom watered at regular intervals. Seedlings were thinned after 1 week to leave only two plants per pot. Whole shoots or different plant parts were harvested as indicated in the legend of figures.

### Cultivation of rice in hydroponics

Rice plants (cv. Nipponbare) were cultivated in hydroponics inside a glasshouse with natural light supplemented with high pressure sodium vapor lamps to ensure a minimum light intensity of 300 μmol m^−2^ s^−1^, and 30°C/25°C day (16 h)/night (8 h) temperature. Seeds were germinated in darkness at 20°C for 3 days and then transferred to meshes floating on a solution containing 0.5 mM CaCl_2_ and 10 μM Na-Fe-EDTA, which was exchanged every third day. After 10 days, homogenous seedlings were transplanted into 60-L tanks filled with half-strength nutrient solution^45^. Ten days later, the nutrient solution was changed to full-strength and exchanged every 10 days. During the whole growing period, the pH value was adjusted to 5.5 every second day. P_i_ starvation was imposed for 10 days before starting P_i_ resupply.

### Cultivation of Physcomitrella patens

*Physcomitrella patens* was grown on Knop medium^46^ solidified with 0.8% agar (A7921, Sigma). Light was provided by fluorescent lamps (60 μmol m^−2^ s^−1^) under a regime of 16 h light and 8 h darkness at constant 20°C. P_i_ treatments were achieved by transferring pre-cultivated plants to fresh Knop solid media containing 1.8 mM KH_2_PO_4_ (+P_i_) or 1.8 mM KCl (-P_i_) for 30 days. At the end of Pi starvation period, part of the plants was resupplied with 1.8 mM KH_2_PO_4_ and harvested after 24 h or 96 h.

### Cultivation of HCT116 cells

Mammalian cells were cultivated as described^23^. Briefly, HCT116 cells were grown in DMEM media supplemented with 10% fetal bovine serum (FBS) and 0.45% glucose in a humidified atmosphere with 5% CO_2_. Phosphate starvation was induced with DMEM without sodium phosphate supplemented with 10% dialyzed FBS. Cells were washed twice in the phosphate-free medium before incubation with DMEM media with or without phosphate. Analysis of InsPs from HCT116 cell lines was performed as previously described^23^.

### Grafting experiment

Seedlings to be grafted were germinated on plates containing half-strength MS (Duchefa), 0.04 mg L^−1^ 6-benzylaminopurine (Sigma), 0.02 mg L^−1^ indole acetic acid (Sigma), and 10 g L^−1^ Difco agar (Becton Dickinson). Five-day-old seedlings were grafted on the plate by the 90-degree blunt end technique with an ultra-fine micro knife (Fine Scientific Tools, USA) without collars. The grafted seedlings remained on the plate for 2 weeks to allow the formation of the graft union. Successfully unified seedlings were transplanted directly to peat-based soil and whole shoots harvested for elemental analysis 2 weeks later.

### RNA isolation and quantitative real-time PCR

Root and shoots tissues were collected by excision and immediately frozen in liquid N_2_. Total RNA was extracted with RNeasy Plant Mini Kit (Macherey-Nagel GmbH & Co KG, Germany). Quantitative reverse transcriptase PCR reactions were conducted with the CFX384TM Real-Time System (Biorad, Germany) and the Go Taq qPCR Master Mix SybrGreen I (Promega) using the primers listed in Supplementary Table 1. *UBQ2* was used as reference gene to normalize relative expression levels of all tested genes. Relative expression was calculated according to published instructions^47^.

### Elemental analysis

Whole shoots were dried at 65°C and digested in concentrated HNO_3_ in polytetrafluoroethylene tubes under pressurized system (UltraCLAVE IV, MLS). Elemental analysis of plant samples from hydroponics or pot experiments was performed by inductively coupled plasma optical emission spectrometry (ICP-OES; iCAP 700, Thermo Fisher Scientific), whereas samples from agar plate-grown plants were analyzed by sector field high-resolution inductively coupled plasma-mass spectrometry (HR-ICP-MS; ELEMENT 2, Thermo Fisher Scientific). Element standards were prepared from certified reference materials from CPI International.

### Titanium dioxide bead extraction and PAGE

InsPs purification and analysis was performed based on a previously established protocol^23, 24,^25. All steps until dilution were performed at 4°C. TiO_2_ beads (Titanium (IV) oxide rutile, Sigma Aldrich) were weighted to 10 mg for each sample and washed once in water and once in 1 M perchloric acid (PA). Liquid N_2_ frozen plant material was homogenized using a pestle and immediately resuspended in 800 μl ice-cold PA. Samples were kept on ice for 10 min with short intermediate vortexing, then centrifuged for 10 min at 20,000×*g* at 4°C using a refrigerated bench top centrifuge. The supernatants were transferred into fresh 1.5 mL tubes and centrifuged again for 10 min at 20,000×*g*. To absorb InsPs onto the beads the supernatants were resuspended in the pre-washed TiO_2_ beads and rotated at 4°C for 30-60 min. Afterwards, beads were pelleted by centrifuging at 8000 *g* for 1 min and washed twice in PA. The supernatants were discarded. To elute inositol polyphosphates, beads were resuspended in 200 μl 10% ammonium hydroxide and then rotated 5 min at room temperature. After centrifuging, the supernatants were transferred into fresh 1.5-mL tubes. The elution process was repeated and the second supernatants were added to the first. Eluted samples were vacuum evaporated at 45°C to dry completely. InsPs were resuspended in 20 μL ultrapure water and separated by 33% polyacrylamide gel electrophoresis and visualized by Toluidine Blue staining, followed by 4′,6-diamidino-2-phenylindole (DAPI) staining.

### ITPK1 *in vitro* kinase and ATP synthase assay

Recombinant *A. thaliana* ITPK1 was purified based on the previously established protocol^48^. The InsP_6_ kinase assay was performed by incubating 10.17 μM enzyme in a reaction mixture containing 5 mM MgCl_2_, 20 mM HEPES (pH 7.5), 1 mM DTT, 5 mM phosphocreatine, 0.33 units creatine kinase, 12.5 mM ATP and 1 mM InsP_6_ (Sichem) at 25°C for 6h. The ability of the enzyme to hydrolyze 5-InsP_7_ was assayed in a reaction mixture containing 3 μg enzyme, 2.5 mM MgCl_2_, 50 mM NaCl, 20 mM HEPES (pH 6.8), 1 mM DTT, 1 mg/mL BSA, 8 mM ADP and 1 mM 5-InsP_7_ at 25°C for 6h. Reactions were separated by 33% polyacrylamide gel electrophoresis and visualized by Toluidine Blue staining.

### NMR-based enzyme assays

Full-length recombinant *A. thaliana* ITPK1 in H_2_O was used in all assays. 0.2-0.8 μM of ITPK1 was incubated in reaction buffer containing 20 mM HEPES pH* 7.0, 50 mM NaCl, 1 mM DTT, 5 mM creatine phosphate, 1 U/ml creatine kinase, 2.5 mM MgCl_2_ (if not indicated otherwise) and 175 μM of [^13^C_6_]InsP_5_ [2-OH], [^13^C_6_]InsP_6_, [^13^C_6_]5-InsP_7_ or [^13^C_6_]1-InsP_7_ in D_2_O. If not indicated otherwise, the reaction buffer also included 2.5 mM ATP or 2.5 mM ADP.

For single time-point analysis of enzyme activity, 2.25-0.375 ng (0.2-0.3 μM) ITPK1 were used. 150 μL reactions were incubated at 25°C (except when 37°C is specified), quenched with 400 μl of 20 mM EDTA (pH* 6.0 in D_2_O) and 11 μL of 5 M NaCl was added for analysis. For real-time monitoring of enzyme activity, 36 ng (0.8 μM) ITPK1 were used. 600 μL reactions were maintained at 25°C in an NMR spectrometer and measured consecutively with 85 sec spectra.

Samples were measured as previously described^49^ on Bruker AV-III spectrometers (Bruker Biospin, Rheinstetten, Germany) using cryogenically cooled 5 mm TCI-triple resonance probe equipped with one-axis self-shielded gradients and operating at 600 MHz for proton nuclei, 151 MHz for carbon nuclei, and 244 MHz for P nuclei. The software to control the spectrometer was topspin 3.5 pl6. Temperature was calibrated using d_4_-methanol and the formula of Findeisen et al.^50^.

### InsPs extraction from seedlings and HPLC analyses

Seedlings were grown vertically on half-strength MS medium supplemented with 1 % sucrose and 7 g L^−1^ Phytagel (P8169, Sigma), pH 5.7 for 12 days (8 h light at 22°C, 16 h darkness at 20°C). Seedlings were transferred to 3 mL half-strength MS liquid media without sucrose and with 625 μM P_i_ (+P) or 5 μM P_i_ (-P). Seedlings were labeled by adding 30 μCi mL^−1^ of [^3^H]-*myo*-inositol (30 to 80 Ci mmol^−1^ and 1 mCi mL^−1^; American Radiolabeled Chemicals) and further cultivated for 5 days. For P_i_-resupply, 620 μM KH_2_PO_4_ was added to the media and the plants were grown for another 6 h before harvest. Afterwards seedlings were washed two times with ultrapure water, frozen in liquid N_2_ and the InsPs were extracted as described previously (Azevedo and Saiardi, 2006). Inositol polyphosphates were resolved by strong anion exchange chromatography HPLC (using a partisphere SAX 4.63 × 125 mm column; HiChrom) at a flow rate of 0.5 mL min^−1^ with a gradient of the following buffers: buffer A (1 mM EDTA) and buffer B (1 mM EDTA and 1.3 M (NH_4_)_2_HPO_4_, pH 3.8, with H_3_PO_4_). The gradient was as follows: 0-2 min, 0 % buffer B; 2-7 min, up to 10 % buffer B; 7-68 min, up to 84 % buffer B; 68-82 min, up to 100 % buffer B; 82-100 min, 100 % buffer B, 100-101 min, down to 0 % buffer B; 101-125 min, 0 % buffer B. Fractions were collected each minute, mixed with scintillation cocktail (Perkin-Elmer; ULTIMA-FLO AP), and analyzed by scintillation counting. To account for differences in fresh weight and extraction efficiencies between samples, values shown are normalized activities based on the total activity of each sample. ‘Total’ activities for normalization were calculated by counting fractions from 26 min (InsP_3_) until the end of the run.

### Analysis of ATP and ADP

Adenosine nucleotides were specifically determined according to Haink and Deussen^51^ with some modifications. 100 mg frozen leaf material from Arabidopsis plants were homogenized in liquid N_2_ and extracted with methanol/chloroform^52^. An aliquot of extracted samples was used for derivatization. Twenty μL of extract was added to 205 μL of a buffer containing 62 mM sodium citrate and 76 mM potassium dihydrogenphosphate for which pH was adjusted to 5.2 with potassium hydroxide. To this mixture, 25 μL chloracetaldehyde (Sigma-Aldrich, Germany) was added and the whole solution was incubated for 40 min at 80°C followed by cooling and centrifugation for 1 min at 14000 rpm. Two blanks containing all reagents except plant extract were used as control. For quantification, external standards (ATP, ADP, AMP) were established with different concentrations. Separation of adenosine nucleotides was performed on a newly developed UPLC-based method using ultra pressure reversed phase chromatography (Acquity H-Class, Waters GmbH, Eschborn, Germany). The UPLC system consisted of a quaternary solvent manager, a sample manager-FTN, a column manager and a fluorescent detector (PDA eλ Detector). The separation was carried out on a C18 reversed phase column (YMC Triart, 1.9 μm, 2.0×100 mm ID, YMC Chromatography, Germany) with a flow rate of 0.6 ml per min and duration of 7 min. The column was heated at 37°C during the whole run. The detection wavelengths were 280 nm for excitation and 410 nm as emission. The gradient was accomplished with two solutions. Eluent A was 5.7 mM tetrabutylammonium bisulfate (TBAS) and 30.5 mM KH_2_PO_4_, pH adjusted to 5.8. Eluent B was a mixture of pure acetonitrile and TBAS in a ratio of 2:1. The column was equilibrated with eluent A (90%) and eluent B (10 %) for at least 30 minutes. The gradient was produced as follow: 90% A and 10% B for 2 min, changed to 40% A and 60% B and kept for 2.3 min, changed to 10% A and 90% B for 1.1 min and reversed to 90% A and 10% B for another 1.6 min.

### Statistical analysis

To analyze the significant differences among multiple groups, one-way analysis of variance followed by Tukey’s test at *P* < 0.05 was adopted. The statistical significance between two groups was assessed by two-tailed Student’s *t*-test. All statistical tests were performed using SigmaPlot 11.0 software.

## Acknowledgments

This work was funded by grants from the Deutsche Forschungsgemeinschaft (DFG, German Research Council) (HE 8362/1-1 to R.F.H.G.; SCHA 1274/4-1, SCHA 1274/5-1, Research Training Group GRK 2064 and under Germany’s Excellence Strategy, EXC-2070-390732324, PhenoRob to G.S.; CIBSS – EXC 2189 – 390939984 as well as JE 572/4-1 to H.J.J.; and LA 4541/1-1, Postdoctoral Research Fellowship, to D.L.) and from the Medical Research Council (MRC) MC_UU_00012/4 to A.S. We thank Annett Bieber, Jacqueline Fuge, Nicole Schäfer and Yudelsy A. Tandron Moya (Leibniz Institute of Plant Genetics and Crop Plant Research) for excellent technical assistance, and Nicolaus von Wirén for critically reading the article. We thank Saikat Bhattacharjee (RCB, India) for providing seeds of published *Arabidopsis thaliana* mutants.

## Author contributions

G.S. and R.F.H.G. conceived the study. D.L., R.K.H., M.F., G.S., and R.F.H.G. designed experiments. E.R., D.L., R.K.H., P.G., V.P., M.F., N.P.L., L.K., R.S., and R.F.H.G. performed experiments. M.-R.H. performed UPLC analysis of ATP and ADP. H.J.J. synthesized various InsP isomers. A.S., D.F., G.S., and R.F.H.G. supervised experimental work. R.F.H.G. and G.S. wrote the manuscript with critical inputs from all authors.

---

**Supplementary Figure 1. Shoot elemental analysis of WT, *itpk1* and recomplemented lines.**

Dry weight of whole shoots (**a**) and shoot concentrations of the macronutrients calcium (**b**), potassium (**c**), magnesium (**d**) and sulfur (**e**), and the micronutrients iron (**f**), and zinc (**g**) of 3-week-old plants grown on peat-substrate. Data represent the mean ± SD (*n* = 8-9 plants). Different letters indicate significant differences according to Tukey’s test (*P* < 0.05).

**Supplementary Figure 2. ITPK1-dependent P overaccumulation in different plant organs.**

Total P_i_ levels in different parts of WT (Col-0) and *itpk1* plants grown on peat-based. Data represent means ± SD (*n* = samples from 5 independent plants). Letters indicate significant differences according to Tukey’s test (*P* < 0.05). Young siliques = green siliques with a length of 0.8 cm to 1.5 cm.

**Supplementary Figure 3. Root phenotype of *itpk1* plants grown in hydroponics and of *pho2-1* grown in agar.**

**a** Phenotype of 5-week-old WT and *itpk1* plants grown in hydroponics with sufficient P_i_. Representative plants are shown. **b** Phenotype of WT and *pho2-1* plants grown in agar plates. Seven-day-old seedlings germinated on half-strength solid MS agar media containing 625 μM P_i_ were transferred to +P (625 μM P_i_) or −P (5 μM P_i_) and grown for additional 7 days.

**Supplementary Figure 4. Expression of PSI genes in *itpk1* plants under different P_i_ conditions.**

Expression analysis of representative P_i_ starvation-induced genes in *itpk1* relative to WT (Col-0). In **a** and **b**, the expression of P_i_ uptake- and signaling-related genes is shown, respectively. Seven-day-old seedlings germinated on half-strength solid MS agar media containing 625 μM P_i_ were transferred to same agar media containing either sufficient (625 μM) or deficient (5 μM) P_i_ levels for 4 days. For P_i_ refeeding, P_i_-deficient plants were transferred back to P_i_-containing media for 6 hours. Data represents means ± SE (*n* = 4 biological replicates). * *P* < 0.05, \*\**P* < 0.01 and \*\*\**P* < 0.001 according to pairwise comparison with Student’s *t*-test (*itpk1* versus Col-0).

**Supplementary Figure 5. P_i_-dependent InsP_7_ and InsP_8_ synthesis is not altered in *itpk2-2* and *itpk4-1* mutant.**

InsP detection in shoots of WT and *itpk2-2* (**a**) and *itpk4-1* plants (**b**) and relative quantification of PAGE signals for *itpk4-1* (**c**). Plants were grown in hydroponics under sufficient P_i_ (+P), after 4 days of P_i_ deficiency (−P) or after resupply of P_i_ to P_i_-deficient plants for 12 h (RS 12h). Data represent means ± SE (*n* = 3 biological replicates).

**Supplementary Figure 6. P_i_-dependent regulation of InsP levels in roots and shoots of rice plants.**

PAGE analysis of rice plants cv. Nipponbare grown in hydroponics under the indicated P_i_ conditions. Shown are representative gels from root and shoot samples.

**Supplementary Figure 7. ITPK1 activity on InsP_5_ [2-OH] and InsP_7_ isomers and control experiments for the kinase activity.**

**a** ITPK1 has no kinase activity on InsP7 isormers. 1-InsP_7_, 5-InsP_7_ or InsP_6_ were incubated with recombinant *Arabidopsis* ITPK1 as indicated in presence of 12.5 mM ATP. InsPs were separated via PAGE and visualized by Toluidine Blue staining. The identity of bands was determined by migration compared to the substrates in absence of enzyme (-). Purified His_8_-MBP tag (MBP) served as negative control for ITPK1. **b-d** Control experiments for NMR analyses. InsP_6_ was incubated with recombinant *Arabidopsis* ITPK1 at 25 °C in the presence of 2.5 mM ATP. Enzymatic activity was determined after 24 h in the presence of varying EDTA concentrations (**b**), after 1.5 h at changing Mg^2+^ concentrations (**c**) and temperature (**d**). The conversion was determined by NMR spectroscopy after quenching with an excess of EDTA. **e-f** 2D^1^H-^13^C-HMBC spectra. Recombinant *Arabidopsis* ITPK1 was incubated with InsP_5_ (**e**) or 1-InsP_7_ (**f**) at 25°C in the presence of an ATP recycling system for 24 h. The reaction mixture analyzed by HSQC NMR spectroscopy. **g** Overview of the reaction shown in (**f**) as analyzed by^31^P NMR spectroscopy after 24 h. A small, unidentified signal potentially reflecting ATP is marked with a question mark.

**Supplementary Figure 8. Dependency of ITPK1 kinase activity on P_i_.**

InsP_6_ was incubated with recombinant *Arabidopsis* ITPK1 at 25°C in the presence of 2.5 mM ATP and the indicated concentrations of P_i_ or its non-metabolizable analog phosphite (Phi). The conversion was determined by NMR spectroscopy after quenching with an excess of EDTA. The experiment was repeated three times.

**Supplementary Figure 9. Recombinant *Arabidopsis* ITPK1 can hydrolyze 5-InsP_7_ in the presence of ADP.**

**a** ^31^P NMR spectroscopy analysis of recombinant *Arabidopsis* ITPK1 incubated with 5-InsP_7_ at 25°C in the presence of ADP. After 24 h the mixture was analyzed by. **b** ^31^P NMR analysis of ATP in ATP synthase reaction buffer. **c** ^31^P NMR spectroscopy analysis of recombinant *Arabidopsis* ITPK1 incubated with ADP without 5-InsP_7_ at 25°C and analyzed after 24 h. A small, unidentified signal potentially reflecting ATP is marked with a question mark.

**Supplementary Figure 10. Effect of P_i_ availability and resupply on shoot ATP levels.**

Concentration of ATP (**a**) and ATP/ADP ratios (**b**) in shoots of Col-0 plants grown in hydroponics with P_i_-sufficient solution (+P), exposed to 4 days of P_i_ starvation (−P) or resupplied with P_i_ for 12 hours (Pi RS 12h). Data represent means ± SE (*n* = 6–7 biological replicates). * *P* < 0.05 and \*\**P* < 0.01 for the indicated pairwise comparisons with Student’s *t*-test.

**Supplementary Figure 11. Shoot ITPK1 function is more determinant for P_i_ accumulation in plants.**

Shoot dry weight and shoot concentrations of the macronutrients potassium (K), calcium (Ca), magnesium (Mg) and the micronutrient iron (Fe) of self-grafted or reciprocally grafted WT (Col-0) and *itpk1*. Plants were grafted on agar plates and left recovering for 2 weeks. Positive grafts were transferred to peat-based substrate for another 2 weeks. Data represent means ± SD (*n* = 5-7 plants). Different letters indicate significant differences according to Tukey’s test (*P* < 0.05). n.s., not significant.

**Supplementary Figure 12. ITPK1-dependent root phenotype in the absence of PHR1 and PHL1.**

Seven-day-old seedlings germinated on half-strength solid MS agar media containing 625 μM P_i_ were transferred to +P (625 μM P_i_) and grown for additional 7 days. Shown are representative images of the indicated mutants grown side-by-side on the same agar plate.

**Supplementary Figure 13. P_i_-dependent transcriptional regulation of InsP-related genes in Col-0 roots.**

Seven-day-old Col-0 seedlings germinated on half-strength solid MS agar media were transferred to the indicated treatments for 4 days. +P, 625 μM P_i_; −P, 5 μM P_i_; Pi RS, P_i_-starved plants were transferred back to P_i_-containing media for 6 hours. Data represent mean ± SE (*n* = 3 biological replicates).

