## Supplementary material for "ITPK1 is an InsP_6_/ADP phosphotransferase that controls systemic phosphate homeostasis in Arabidopsis": Suppl. Information

### Supplementary Information – Riemer et al.

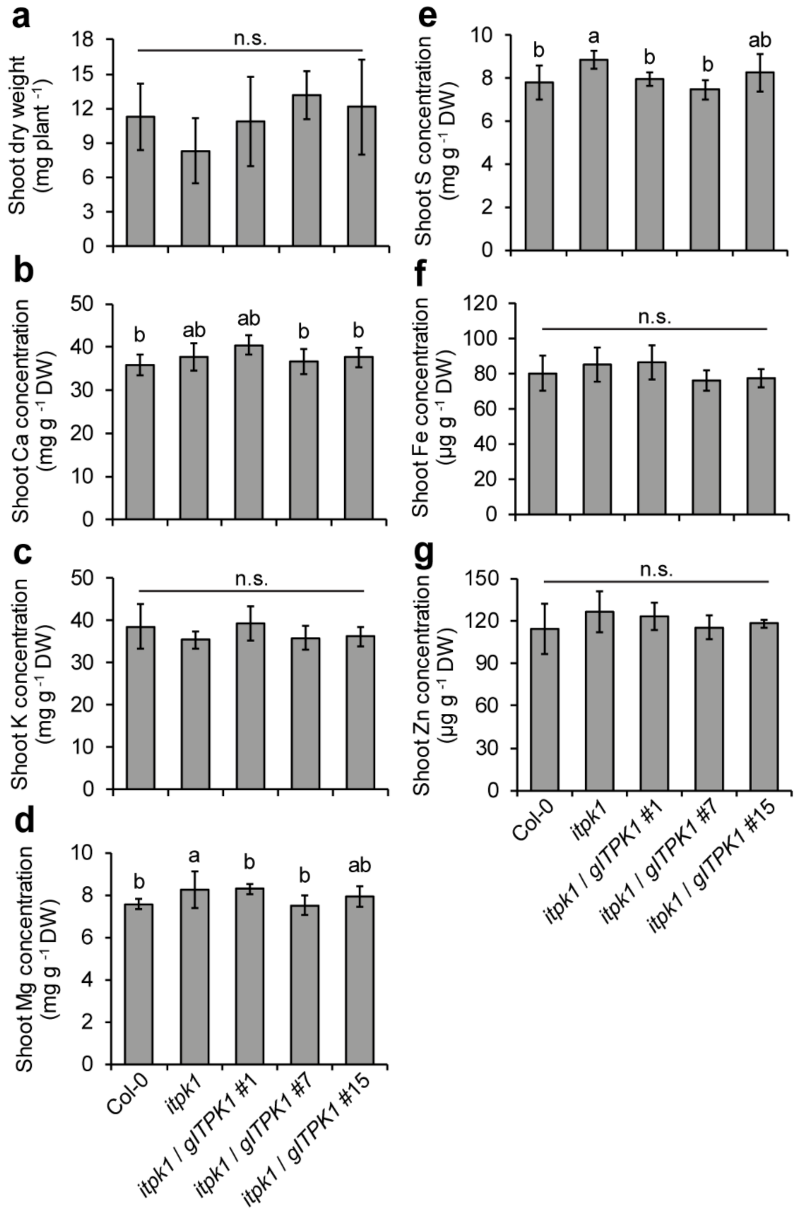


**Supplementary Figure 1. Shoot elemental analysis of WT, *itpk1* and recomplemented lines.** Dry weight of whole shoots (**a**) and shoot concentrations of the macronutrients calcium (**b**), potassium (**c**), magnesium (**d**) and sulfur (**e**), and the micronutrients iron (**f**), and zinc (**g**) of 3-week-old plants grown on peat-substrate. Data represent the mean ± SD ($n$ = 8-9 plants). Different letters indicate significant differences according to Tukey's test ($P < 0.05$).

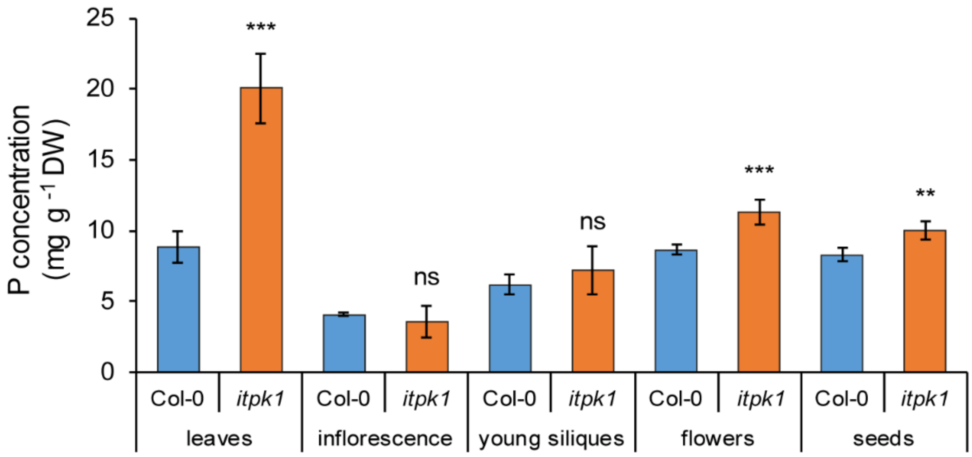


**Supplementary Figure 2. ITPK1-dependent P overaccumulation in different plant organs.**
Total $P_i$ levels in different parts of WT (Col-0) and *itpk1* plants grown on peat-based. Data represent means ± SD (*n* = samples from 5 independent plants). Letters indicate significant differences according to Tukey's test ($P < 0.05$). Young siliques = green siliques with a length of 0.8 cm to 1.5 cm.

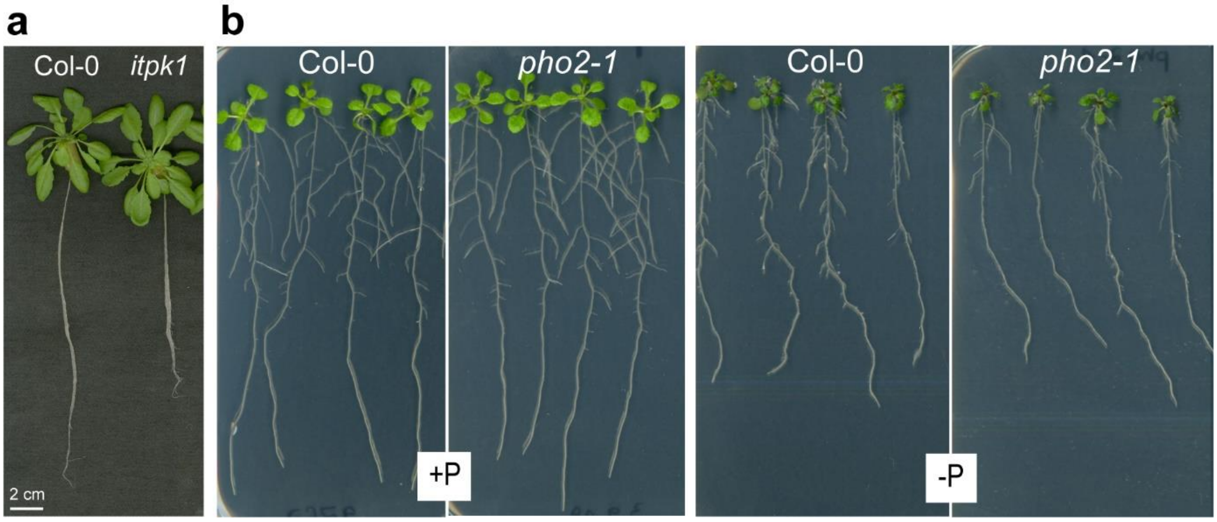


**Supplementary Figure 3. Root phenotype of *itpk1* plants grown in hydroponics and of *pho2-1* grown in agar.**
**a** Phenotype of 5-week-old WT and *itpk1* plants grown in hydroponics with sufficient $P_i$. Representative plants are shown. **b** Phenotype of WT and *pho2-1* plants grown in agar plates. Seven-day-old seedlings germinated on half-strength solid MS agar media containing 625 µM $P_i$ were transferred to +P (625 µM $P_i$) or -P (5 µM $P_i$) and grown for additional 7 days.

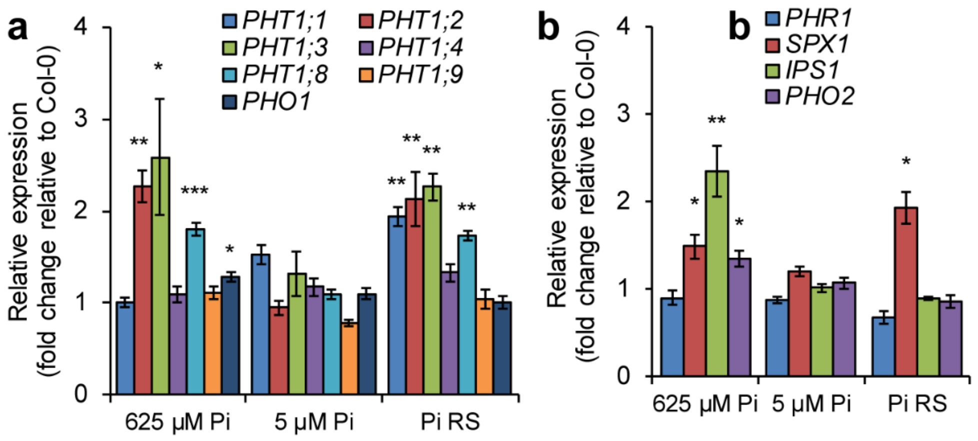


**Supplementary Figure 4. Expression of PSI genes in *itpk1* plants under different $P_i$ conditions.** Expression analysis of representative $P_i$ starvation-induced genes in *itpk1* relative to WT (Col-0). In **a** and **b**, the expression of $P_i$ uptake- and signaling-related genes is shown, respectively. Seven-day-old seedlings germinated on half-strength solid MS agar media containing 625 µM $P_i$ were transferred to same agar media containing either sufficient (625 µM) or deficient (5 µM) $P_i$ levels for 4 days. For $P_i$ refeeding, $P_i$-deficient plants were transferred back to $P_i$-containing media for 6 hours. Data represents means ± SE ($n$ = 4 biological replicates). * $P < 0.05$, **$P < 0.01$ and ***$P < 0.001$ according to pairwise comparison with Student's $t$-test (*itpk1* versus Col-0).

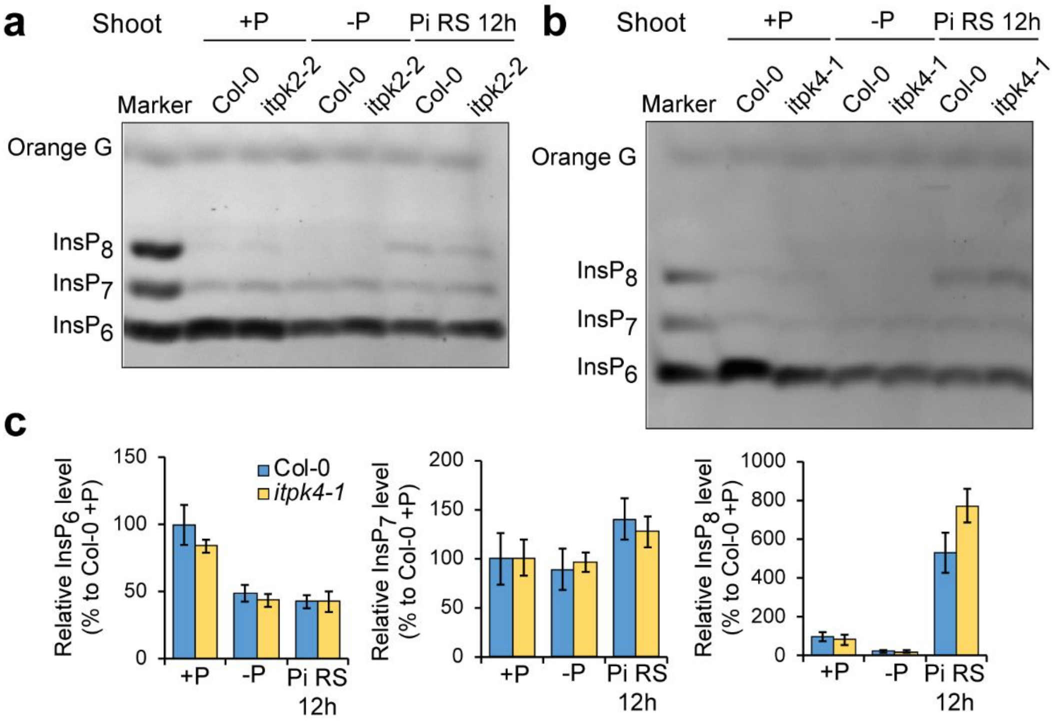


**Supplementary Figure 5. $P_i$-dependent $InsP_7$ and $InsP_8$ synthesis is not altered in *itpk2-2* and *itpk4-1* mutant.**
InsP detection in shoots of WT and *itpk2-2* (**a**) and *itpk4-1* plants (**b**) and relative quantification of PAGE signals for *itpk4-1* (**c**). Plants were grown in hydroponics under sufficient $P_i$ (+P), after 4 days of $P_i$ deficiency (-P) or after resupply of $P_i$ to $P_i$-deficient plants for 12 h (RS 12h). Data represent means ± SE ($n$ = 3 biological replicates).

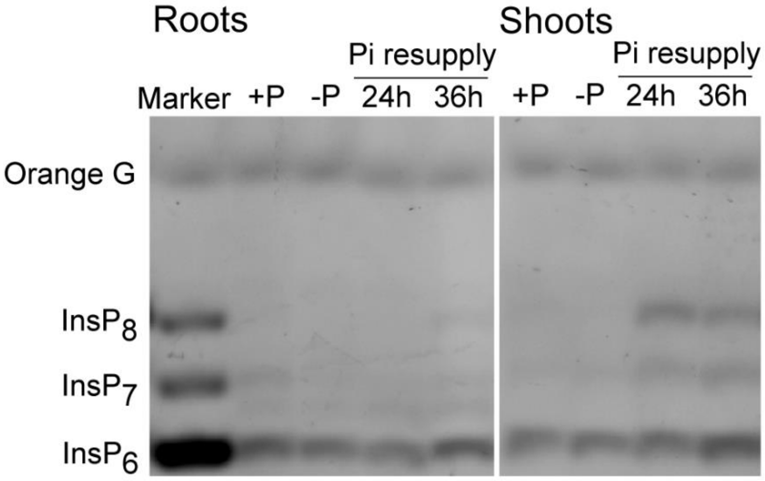


**Supplementary Figure 6. $P_i$-dependent regulation of InsP levels in roots and shoots of rice plants.** PAGE analysis of rice plants cv. Nipponbare grown in hydroponics under the indicated $P_i$ conditions. Shown are representative gels from root and shoot samples.

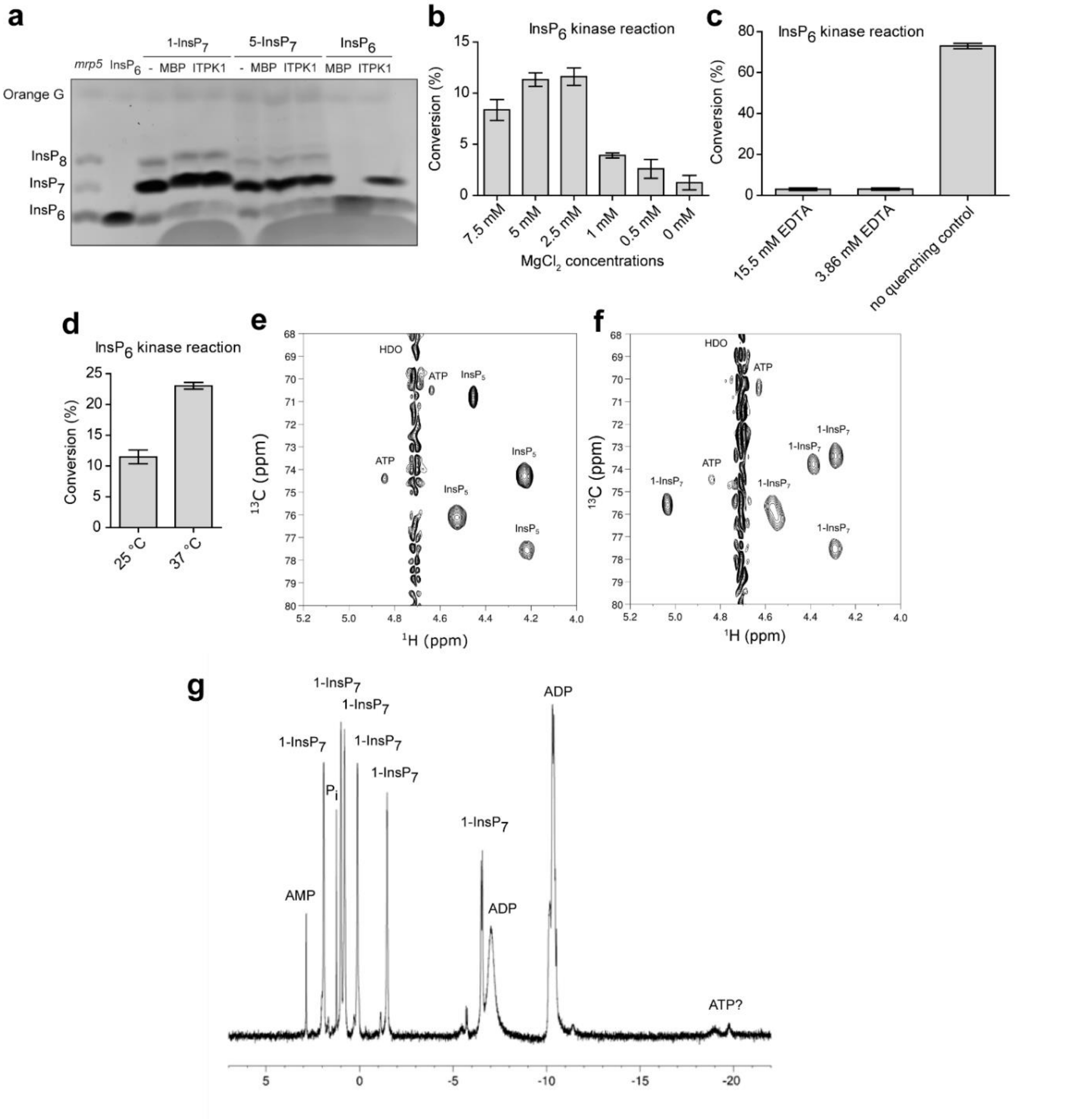


**Supplementary Figure 7. ITPK1 activity on $InsP_5$ [2-OH] and $InsP_7$ isomers and control experiments for the kinase activity.**

**a** ITPK1 has no kinase activity on InsP7 isormers. 1-$InsP_7$, 5-$InsP_7$ or $InsP_6$ were incubated with recombinant *Arabidopsis* ITPK1 as indicated in presence of 12.5 mM ATP. InsPs were separated via PAGE and visualized by Toluidine Blue staining. The identity of bands was determined by migration compared to the substrates in absence of enzyme (**-**). Purified $His_8$-MBP tag (MBP) served as negative control for ITPK1. **b-d** Control experiments for NMR analyses. $InsP_6$ was incubated with recombinant *Arabidopsis* ITPK1 at 25 °C in the presence of 2.5 mM ATP. Enzymatic activity was determined after 24 h in the presence of varying EDTA concentrations (**b**), after 1.5 h at changing $Mg^{2+}$ concentrations (**c**) and temperature (**d**). The conversion was determined by NMR spectroscopy after quenching with an excess of EDTA. **e-f** 2D $^{1}H$-$^{13}C$-HMBC spectra. Recombinant *Arabidopsis* ITPK1 was incubated with $InsP_5$ (**e**) or 1-$InsP_7$ (**f**) at 25°C in the presence of an ATP recycling system for 24 h. The reaction mixture analyzed by HSQC NMR spectroscopy. **g** Overview of the reaction shown in (**f**) as analyzed by $^{31}P$ NMR spectroscopy after 24 h. A small, unidentified signal potentially reflecting ATP is marked with a question mark.

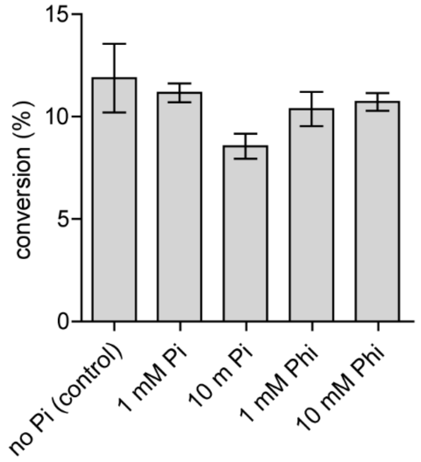


**Supplementary Figure 8. Dependency of ITPK1 kinase activity on $P_i$.**

$InsP_6$ was incubated with recombinant *Arabidopsis* ITPK1 at 25°C in the presence of 2.5 mM ATP and the indicated concentrations of $P_i$ or its non-metabolizable analog phosphite (Phi). The conversion was determined by NMR spectroscopy after quenching with an excess of EDTA. The experiment was repeated three times.

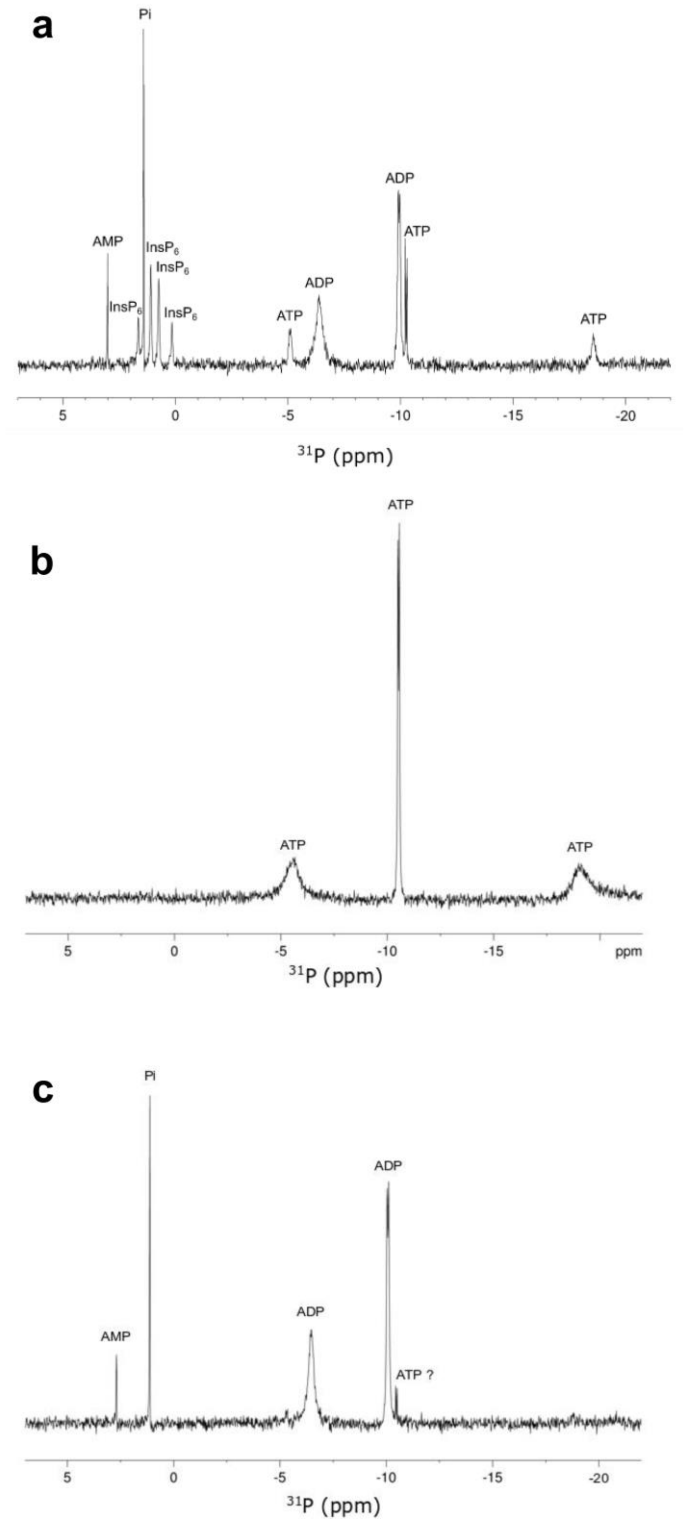


**Supplementary Figure 9. Recombinant *Arabidopsis* ITPK1 can hydrolyze 5-InsP$_7$ in the presence of ADP.**

**a** $^{31}$P NMR spectroscopy analysis of recombinant *Arabidopsis* ITPK1 incubated with 5-InsP$_7$ at 25°C in the presence of ADP. After 24 h the mixture was analyzed by. **b** $^{31}$P NMR analysis of ATP in ATP synthase reaction buffer. **c** $^{31}$P NMR spectroscopy analysis of recombinant *Arabidopsis* ITPK1 incubated with ADP without 5-InsP$_7$ at 25°C and analyzed after 24 h. A small, unidentified signal potentially reflecting ATP is marked with a question mark.

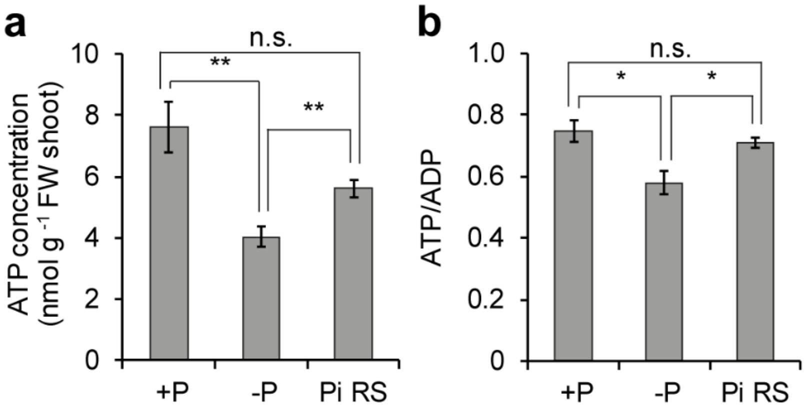


**Supplementary Figure 10. Effect of $P_i$ availability and resupply on shoot ATP levels.**
Concentration of ATP (**a**) and ATP/ADP ratios (**b**) in shoots of Col-0 plants grown in hydroponics with $P_i$-sufficient solution (+P), exposed to 4 days of $P_i$ starvation (-P) or resupplied with $P_i$ for 12 hours (Pi RS 12h). Data represent means ± SE ($n$ = 6–7 biological replicates). * $P < 0.05$ and **$P < 0.01$ for the indicated pairwise comparisons with Student's *t*-test.

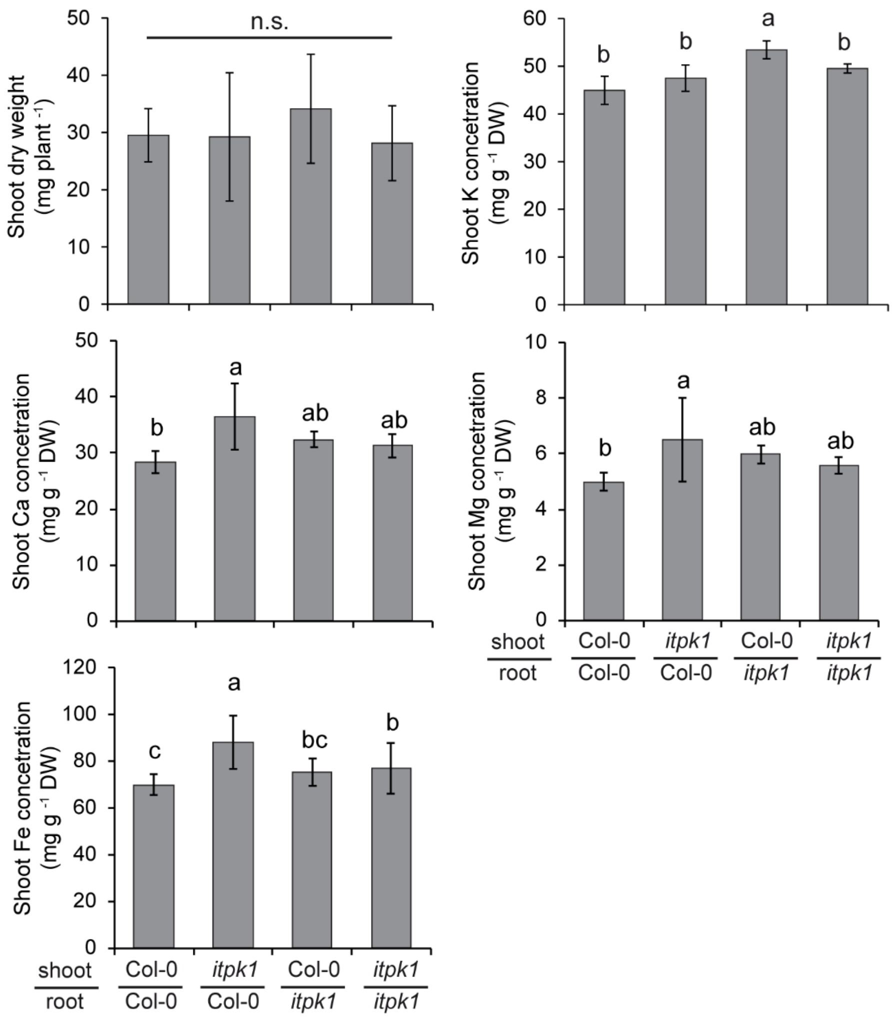


**Supplementary Figure 11. Shoot ITPK1 function is more determinant for $P_i$ accumulation in plants.**

Shoot dry weight and shoot concentrations of the macronutrients potassium (K), calcium (Ca), magnesium (Mg) and the micronutrient iron (Fe) of self-grafted or reciprocally grafted WT (Col-0) and *itpk1*. Plants were grafted on agar plates and left recovering for 2 weeks. Positive grafts were transferred to peat-based substrate for another 2 weeks. Data represent means ± SD ($n$ = 5-7 plants). Different letters indicate significant differences according to Tukey's test ($P < 0.05$). n.s., not significant.

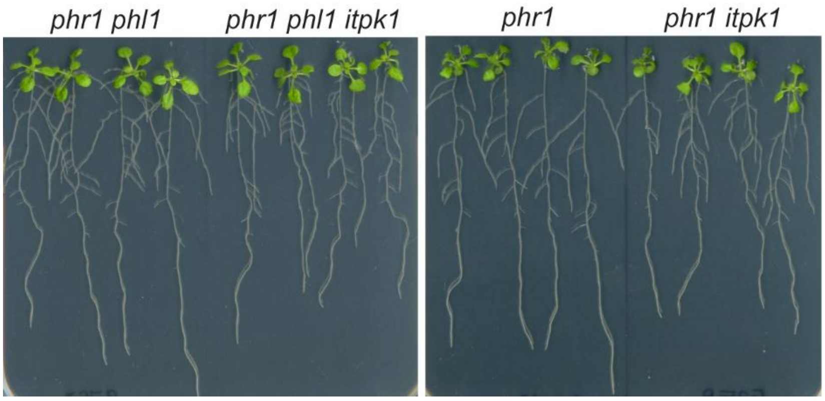


**Supplementary Figure 12. ITPK1-dependent root phenotype in the absence of PHR1 and PHL1.** Seven-day-old seedlings germinated on half-strength solid MS agar media containing 625 µM $P_i$ were transferred to +P (625 µM $P_i$) and grown for additional 7 days. Shown are representative images of the indicated mutants grown side-by-side on the same agar plate.

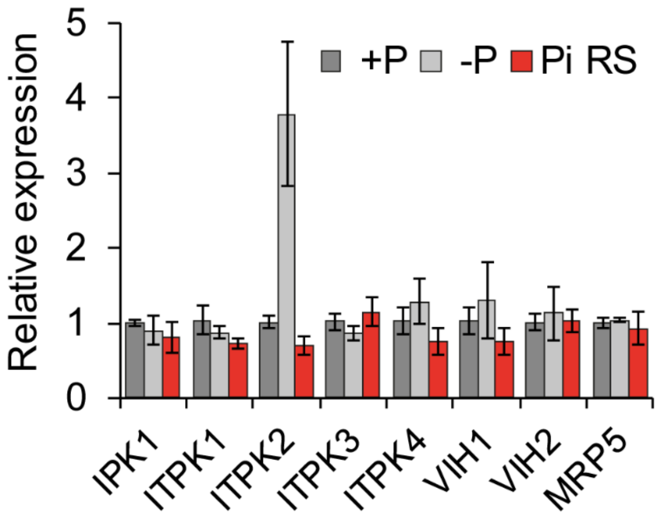


**Supplementary Figure 13. $P_i$-dependent transcriptional regulation of InsP-related genes in Col-0 roots.**

Seven-day-old Col-0 seedlings germinated on half-strength solid MS agar media were transferred to the indicated treatments for 4 days. +P, 625 µM $P_i$; -P, 5 µM $P_i$; Pi RS, $P_i$-starved plants were transferred back to $P_i$-containing media for 6 hours. Data represent mean ± SE ($n$ = 3 biological replicates).

**Supplementary Table 1. List of primers used in this study.**

| AGI ID | Gene name | Primer sequence |
|---|---|---|
| **Primers used for qPCR analysis** | | |
| AT2G36170 | *UBQ2* | F: 5’-CCAAGATCCAGGACAAAGAAGGA-3’<br>R: 5’-TGGAGACGAGCATAACACTTG-3’ |
| AT5G43350 | *PHT1;1* | F: 5’-AGGCGATCACGTTGCTTACA-3’<br>R: 5’- TCTCTGGAGAGTTGAGGAGAGAC-3’ |
| AT5G43370 | *PHT1;2* | F: 5’-CCATTAGCGCACAACGGAAAG-3’<br>R: 5’-GAAACCCATACCGGCGATGA-3’ |
| AT5G43360 | *PHT1;3* | F: 5’-GCTTTCATCGCGGCAGTGTT-3’<br>R: 5’-TGAGGAGGCGTTGATAGAAACC-3’ |
| AT2G38940 | *PHT1;4* | F: 5’-AGCCTTTGTCTCTGCGGTTT-3’<br>R: 5’-CGTGGATCCCAAGGCATCAT-3’ |
| AT1G20860 | *PHT1;8* | F: 5’-AGAGAAGTGGCGGTGGTTTG-3’<br>R: 5’-TCTTGCGGTTTCAGGCATCA-3’ |
| AT1G76430 | *PHT1;9* | F: 5’-TTCGGAGAAGACGAACGTGG-3’<br>R: 5’-GTATCTGGCGGTTTCAGGCA-3’ |
| AT3G23430 | *PHO1* | F: 5’-GACTTACAGCTCGTTGAATATGATAGC-3’<br>R: 5’-CGATCTCTTTACGACTTTGAGATACG-3’ |
| AT4G28610 | *PHR1* | F: 5’-TGTGGAATTGCGACCTGTTA-3’<br>R: 5’-GCTCTTTCACTACCGCCAAG-3’ |
| AT5G20150 | *SPX1* | F: 5’-GTTGATTTCCATGGAGAAATGG-3’<br>R: 5’-GGTAAACGCATGAGATCACCAG-3’ |
| AT3G09922 | *IPS1* | F: 5’-TCCCTCTAGAAATTGGGCAAC-3’<br>R: 5’-GGGAGTGGGTACAACCCAAA-3’ |
| AT2G33770 | *PHO2* | F: 5’-TTGCACCATGTGAAATTTGG-3’<br>R: 5’-AGACCCGTTTCCTGATGGTT-3’ |
| AT5G42810 | *IPK1* | F: 5’-TTACAGAGGCGAAGGTGGTG-3’<br>R: 5’-AGGACCCAAGAGTGGGATGA-3’ |
| AT5G16760 | *ITPK1* | F: 5’-ATTGGGACGTCGAAAGGGTC-3’<br>R: 5’-CTCAGTCAACACAGGCTCGT-3’ |
| AT4G33770 | *ITPK2* | F: 5’-ATTTAGGGGTTTCCTGCGGT-3’<br>R: 5’-GTTGAAACTGGAACGGCGTC-3’ |
| AT4G08170 | *ITPK3* | F: 5’-CTTTCAGAGCAGGGTCCGTT-3’<br>R: 5’-ACACCAACGCGTCCATTACT-3’ |
| AT2G43980 | *ITPK4* | F: 5’-TGTGAAACCACAGGTTGCCT-3’<br>R: 5’-TCGATTTCTTGACCGCGTGA-3’ |
| AT5G15070 | *VIH1* | F: 5’-AGCTACGCTGTGTCATTGCT-3’<br>R: 5’-CACCCTCTTCAAGAACGGCT-3’ |
| AT3G01310 | *VIH2* | F: 5’-AGCATTGAAATGGAAGCGGC-3’<br>R: 5’-GTTGGGAGGAAGTCCAGCTC-3’ |
| AT1G04120 | *MRP5* | F: 5’-ATAGAAGATTTCCGCCCGCC-3’<br>R: 5’-TTTCCACCCGGAAACACACA-3’ |
| **Primers used for genotyping in F1, F2 and F3** | | |
| itpk1_LP | 5’-ACCAATATTCGATTCCACACG-3’ | |
| itpk1_RP | 5’-CCATGTCCCAGAAGAACTCAG-3’ | |
| SAIL_LB3 | 5’-TAGCATCTGAATTTCATAACCAATCTCGATACAC-3’ | |
| phr1_LP | 5’-GAGAGACCTCACACGCACTTC-3’ | |
| phr1_RP | 5’-CTTTCTGGCGAACCTGTAGTG-3’ | |
| phl1_LP | 5’-GTGGAGACGTTTCTGCACTTC-3’ | |
| phl1_RP | 5’-TCCCACAATCCAAATTCAGAG-3’ | |
| LBb1.3 | 5’-ATTTTGCCGATTTCGGAAC-3’ | |
